# Electroconvulsive Therapy Induces Dopaminergic Axon Regeneration, Corticostriatal Remodeling, and Restoration of Motor Function in Parkinsonian Mice

**DOI:** 10.64898/2025.12.05.692581

**Authors:** Anika Frank, Se Joon Choi, Jonas Bendig, Anna S. Monzel, Adrienne C. Ferguson, Siham Boumhaouad, Martin Picard, Eugene V. Mosharov, David Sulzer

**Affiliations:** Department of Psychiatry, Division of Molecular Therapeutics, Columbia University Irving Medical Center (CUIMC), New York, NY, USA; New York State Psychiatric Institute, RFMH, New York, NY, USA; Department of Neurology, H. Houston Merritt Center, Columbia Translational Neuroscience Initiative, CUIMC, New York, NY, USA; Robert N. Butler Columbia Aging Center, Columbia University Mailman School of Public Health, New York, NY, USA; Departments of Neurology and Pharmacology, CUIMC, New York, NY, USA

## Abstract

Parkinson’s disease (PD) is characterized by the progressive degeneration of midbrain dopaminergic neurons with loss of axonal dopamine neurotransmission in the dorsal striatum, leading to striatal circuit dysfunction and debilitating motor symptoms. Current therapies provide symptomatic relief but do not restore lost neuronal function. Decades of clinical observations have reported the unexpected observation that electroconvulsive therapy (ECT), a standard treatment for refractory neuropsychiatric disorders, can incidentally alleviate motor symptoms in PD patients, yet the underlying mechanisms remain unknown. Here, we report that in the unilateral 6-hydroxydopamine (6-OHDA) mouse model of PD, two weeks of repeated ECT produced robust and sustained motor recovery, with improvements in locomotion and sensorimotor asymmetry persisting for at least 30 days post-treatment. Remarkably, ECT induced dopaminergic axonal sprouting from surviving dopaminergic neurons with cell bodies located at the substantia nigra-ventral tegmental border, leading to a partial recovery of striatal dopaminergic axonal reinnervation. In striatal direct pathway spiny projection neurons (dSPNs), which exhibit pathological hyperexcitability and spine loss following dopamine depletion, ECT normalized both corticostriatal synaptic responses and spine density. Consistently, ECT upregulated gene transcripts involved in cytoskeletal remodeling while downregulating those associated with glutamatergic signaling and neuronal excitability. These changes were accompanied by a coordinated transcriptional shift toward enhanced mitochondrial anabolic capacity and energy production, including increased expression of genes involved in ATP and Coenzyme Q biosynthesis. Together, these findings demonstrate that ECT can partially restore basal ganglia circuitry following dopamine depletion and provide a basis for further study of its potential as a noninvasive, disease-modifying intervention for PD.

## INTRODUCTION

Parkinson’s disease (PD) is marked by the progressive loss of dopaminergic neurons in the substantia nigra (SN), leading to cardinal motor symptoms such as rigidity, bradykinesia, and tremor. Additional brain regions and neurotransmitter systems are affected, resulting in a range of non-motor neuropsychiatric symptoms including depression, anxiety, and dementia^1^.

The striatum, a key hub of motor function that receives massive dopaminergic input from the SN along with glutamatergic afferents from the thalamus, cortex, and other brain regions^2,3,4^, is profoundly affected in PD. The loss of striatal dopamine axons reduces excitatory signaling via D1 dopamine receptors on direct pathway spiny projection neurons (dSPNs) and inhibitory signaling via D2 dopamine receptors on indirect pathway SPNs (iSPNs), resulting in hypoactivity of dSPNs and hyperactivity of iSPNs. This imbalance in dopaminergic signaling, accompanied by substantial SPN dendritic atrophy and dendritic spine loss, disrupts basal ganglia output producing multiple PD motor deficits, including bradykinesia, rigidity, and tremor^3,5^.

In PD patients, dopamine replacement therapy by its precursor levodopa (L-3,4-dihydroxyphenylalanine, L-DOPA) is the primary treatment for motor symptoms. The efficacy of L-DOPA can persist for several days after cessation of the drug (“long duration response”), suggesting that the treatment underlies a long-lasting normalization of synaptic responses and structures^6^. Consistently, L-DOPA has been shown to partially normalize SPN activity and morphology^7,8^. However, as dopaminergic degeneration progresses, chronic administration of higher doses of L-DOPA is required, leading to motor complications, including involuntary movements (dyskinesia) and motor fluctuations(“on-off” phenomenon)^3^. These observations highlight the critical need for alternative therapeutic strategies that can achieve targeted, long-term modulation of pathological neuronal circuits without the complication profile of associated with chronic L-DOPA use^9,10^.

Given its ability to induce neuroplastic changes^11,12^, electroconvulsive therapy (ECT), may provide such a therapeutic avenue. Introduced as a psychiatric treatment in the 1930s, ECT is now largely considered to provide the most effective therapy for patients with drug-resistant depression and psychosis^13^. While ECT use has declined with the introduction of modern antidepressants, innovations such as unilateral ultra-brief pulse protocols and a growing need for effective therapies for drug-resistant depression have led to a resurgence^14^ and in the U.S. it is presently used in ∼10,000 patients (∼ 70,000 treatments) annually^15^. This modified ECT is administered under general anesthesia and muscle relaxation and involves a brief electrical stimulation lasting up to 8 seconds to elicit a controlled seizure lasting up to 60 seconds^16^.

Depression is highly prevalent in PD patients^17^, and ECT has therefore occasionally been employed to treat these comorbid mood symptoms. ECT is generally well tolerated with most of its side effects, such as headache, muscle soreness, or short-term memory disturbances^13^, being transient. Moreover, its demonstrated effectiveness in older individuals makes it an appropriate therapeutic option for PD patients^18^. A meta-analysis of fourteen clinical studies suggests that ECT alleviates depression in PD, but surprisingly, also improves PD motor symptoms, even in patients without psychiatric comorbidities^19^. Most of these findings are based on case reports and small clinical trials, with only a few studies employing controlled blinding. One study in non-depressed, non-demented PD patients demonstrated significant motor improvements post-ECT, including a nearly 50% reduction in motor symptom severity scores with improvements in walking time and step count, symptoms that are often resistant to deep brain stimulation, a current clinical standard treatment in advanced PD^20,21^. A double-blind study found that ECT-treated patients with L-DOPA “on-off phenomenon” experienced a 34% longer duration of “on” phases (periods of good motor function during the day), compared to a 0.5% increase in sham-treated controls^22^.

Despite these reports, ECT is largely unexplored as a therapy for PD, primarily because of a lack of mechanistic insights that could explain its benefits. Notably, even in the treatment of depression, where ECT has a long history and wide clinical use, its biological effects remain incompletely understood. Several neuroplasticity-based hypotheses have been suggested^23^, including the release of brain-derived neurotrophic factor (BDNF)^24,25^, induction of hippocampal neurogenesis^12^ and an increased number of GABAergic interneurons in the medial striatum^26^. However, to our knowledge, these mechanisms have not been directly tested in the context of PD.

Here, we report that ECT produces prominent and long-lasting behavioral recovery in a partial dopamine-lesion mouse model of PD. This recovery is accompanied by ECT-dependent reinnervation of the dorsal motor striatum by dopaminergic axons from surviving neurons, along with normalization of electrophysiological and morphological properties of the striatal dSPNs, consistent with known effects of dopamine replacement on SPNs. Single-nucleus RNA sequencing of dSPNs revealed an ECT-elicited homeostatic transcriptional program marked by upregulation of genes involved in mitochondrial function and cytoskeletal remodeling, and downregulation of gene networks associated with glutamatergic receptor overactivation and heightened intrinsic excitability. Together, these findings position ECT as a noninvasive, neurorestorative intervention that repairs basal ganglia circuitry through enhanced bioenergetics, axonal sprouting, and synaptic regeneration.

## RESULTS

### ECT provides rapid and long-lasting improvement of parkinsonian motor behaviors

To assess the therapeutic effects of ECT in PD, we used the unilateral partial 6-hydroxydopamine (6-OHDA) lesion model in which injection of the toxin into the dorsal striatum selectively targets dopaminergic axons and cell bodies of the substantia nigra (SN) in one hemisphere, while sparing dopaminergic neurons in the ventral tegmental area and their projections to the ventral striatum^27^. Three weeks after lesioning, animals underwent baseline behavioral testing before receiving the intervention. Treatment consisted of either whole-brain ECT (1 second per day, 5 sessions per week for two weeks) or sham treatment (isoflurane anesthesia only). Post-intervention behavioral assessments were performed 3 days and 4 weeks after the final ECT treatment (**Fig. 1a**).

**Figure 1.**
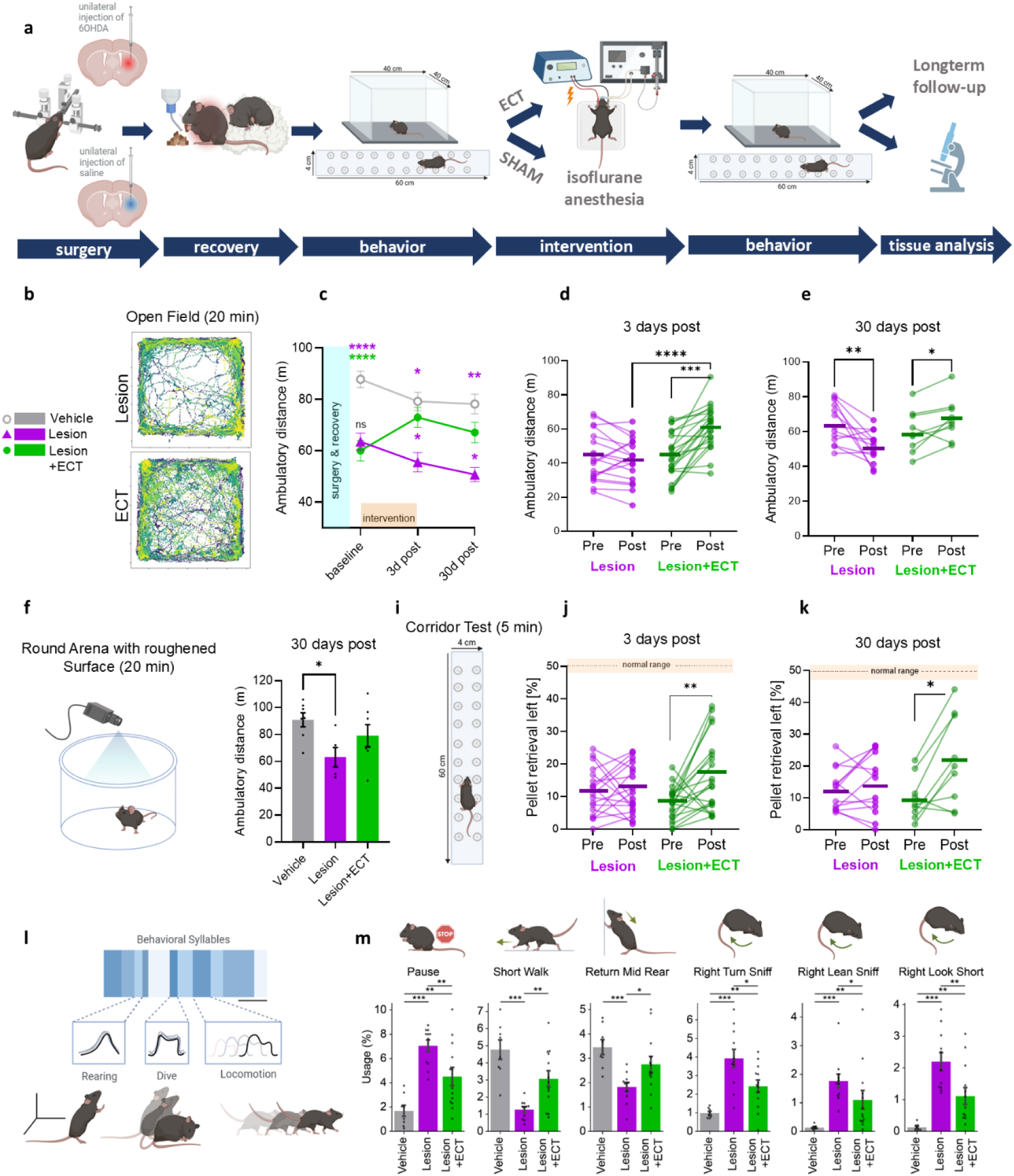
ECT provides rapid and long-lasting improvement of parkinsonian motor behaviors. (**a**) Schematic overview of the experimental design. The PD phenotype was induced by unilateral injection of 6-OHDA into the dorsal striatum of wild-type mice. Behavioral baseline data was obtained 3 weeks after 6-OHDA injection and mice were randomized into either sham treatment (1 min isoflurane anesthesia) or ECT (1 sec of 50 mA current pulse applied via ear clips under anesthesia). Treatment groups are Vehicle (saline), Lesion (6-OHDA + sham treatment), and Lesion+ECT (6-OHDA + verum treatment). ECT or sham sessions were performed 10 times over 2 weeks, and behavioral tests were repeated 3 or 30 days after the last treatment. Mice were then euthanized for electrophysiology, immunofluorescence, or RNA-seq analysis. (**b**) Representative open field movement traces from Lesion and Lesion+ECT mice. (**c**) Average changes in total ambulatory distance measured before and after the intervention (two-way ANOVA with post-hoc Tukey’s HSD; Vehicle (n=8), Lesion (n=12), and Lesion+ECT(n=10)). (**d-f**) Effect of dopamine lesion and ECT on individual animals’ mobility; ECT significantly increased total distance traveled 3 days after the intervention (d, Lesion (n=19), and Lesion+ECT(n=22)) and prevented a decline in locomotion activity at 30 days (e, Lesion (n=13), and Lesion+ECT(n=9)). (**f**) Ambulatory distance was assessed in a novel environment using a circular arena with a roughly sanded surface. Lesioned mice that received sham treatment exhibited reduced activity, whereas ECT-treated mice displayed locomotor behavior comparable to vehicle animals (Vehicle (n=7), Lesion (n=5), and Lesion+ ECT (n=7)). (**i-k**) Corridor test setup (i) and individual animal performance at 3 (j) and 30 (k) days post-ECT. Whereas unlesioned mice retrieved food pellets equally from both sides (shown as dotted line at 50%), unilateral 6-OHDA lesion caused retrieval asymmetry, which was partially restored by ECT treatment. Group sizes are the same as in panel (d) and (e). (**l**) Unsupervised Motion Sequencing (MoSeq) performed 3 days after ECT revealed 30 distinct movement syllables that were changed by the lesion, 18 of which were significantly restored by ECT (see Supplementary Fig. 1). (**m**) Examples of behavior syllables with their usage frequency (% of total time) during a 20-minute session (Kruskal-Wallis with post-hoc Dunn; Vehicle (n=8), Lesion (n=11), and Lesion+ECT(n=13)). One-way ANOVA was used to compare means across multiple independent groups with post-hoc Tukey’s HSD unless indicated otherwise. Paired t-tests were applied for repeated measurements within the same group, while unpaired t-tests were used for comparisons between two independent groups. * p < 0.05, ** p < 0.01, *** p < 0.001, **** p<0.0001. Cartoons were prepared with BioRender.com. Panels (f) and (i) are adapted from Lin et al.^31^

Consistent with previous reports, unilaterally 6-OHDA-lesioned mice displayed motor deficits including contralateral forelimb akinesia, spontaneous ipsilateral rotations, and decreased horizontal and vertical activity^28^. On the open field test, lesioned mice showed greatly reduced ambulatory distance compared to saline-injected controls (**Fig. 1 c**). ECT-treatment significantly ameliorated the decline in mobility both 3 days and 4 weeks post-intervention, while sham-treated dopamine-lesioned mice exhibited a progressive decline in mobility (**Fig. 1d-e**). To ensure our assessment of locomotor activity was independent of effects of habituation to a standard open-field box, we measured the total distance traveled in a novel environment, a round arena with a sanded surface, in contrast to the square, smooth-surfaced arena used previously. In the novel environment, ECT-treated mice showed locomotor activity comparable to the Vehicle group, while the Lesion group remained less active (**Fig. 1f**).

We then employed the corridor test to assess sensorimotor asymmetry^29^. At baseline, 6-OHDA-lesioned mice retrieved and consumed only ∼10% of sugar pellets from the contralateral (left) side of their body, compared to ∼50% retrievals and explorations of vehicle controls, indicating a strong lateralized motor deficit. ECT treatment partially restored retrieval behavior on the affected side, after both 3 days and 4 weeks post-intervention, while sham-treated PD mice maintained their lateralized deficit (**Fig. 1i-k**), further supporting a beneficial impact of ECT on motor function.

To further dissect behavioral changes with greater granularity, we performed motion sequencing (MoSeq) analysis. At 3 days post-treatment, MoSeq identified 30 distinct behavioral ‘syllables’ – discrete, stereotyped units of movement – with different usage frequency in the Lesion group compared to Vehicle controls (minimum cumulative syllable usage of 3 % across groups, **Fig. 1l-m, Supplementary Fig. 1**). Of these, 13 syllables were partially restored and 5 were fully rescued by ECT. These syllables were associated with increased rearing behavior and high-velocity dashes, behaviors typically reduced in 6-OHDA parkinsonian mice^30^. Moreover, ECT-treated mice exhibited less movement fragmentation, characterized by fewer pauses during ambulatory phases, and a notable reduction in spontaneous rotational behavior (**Fig. 1m**). Similar behavioral syllables showed recovery when mice were tested 4 weeks after ECT (**Supplementary Fig. 1**), confirming that the procedure produced long-lasting improvement in parkinsonian features.

### ECT promotes dopamine axon sprouting in the striatum

Similar improvement of 6-OHDA induced akinesia and lesion-induced ipsilateral bias in the corridor test has been shown by dopamine replacement therapy^32,33^. To investigate the structural correlates of the observed behavioral recovery, we assessed whether ECT promotes dopaminergic axon sprouting and reinnervation within the dorsal striatum of 6-OHDA-lesioned mice.

Tyrosine hydroxylase (TH) immunolabeling on coronal brain sections at approximately +0.5 mm anterior to bregma showed the expected 6-OHDA-mediated lesion of TH-positive fibers in both dorsal and ventral striatum (**Fig. 2a**). Strikingly, ECT-treated mice exhibited a significant increase in striatal TH+ fiber density, corresponding to a ∼2.5-fold increase relative to the Lesion group (**Fig. 2b**). The dorsolateral striatum displayed distinct morphological features of active axonal remodeling, including thin, varicose processes with irregular trajectories. Many of these fibers exhibited terminal structures resembling growth cones, consistent with axonal sprouting (**Fig. 2c-d**). In contrast, lesioned animals exhibited comparatively sparse and less complex TH^+^ processes in the same region, with fewer growth cone-like structure than those seen in ECT-treated animals.

**Figure 2.**
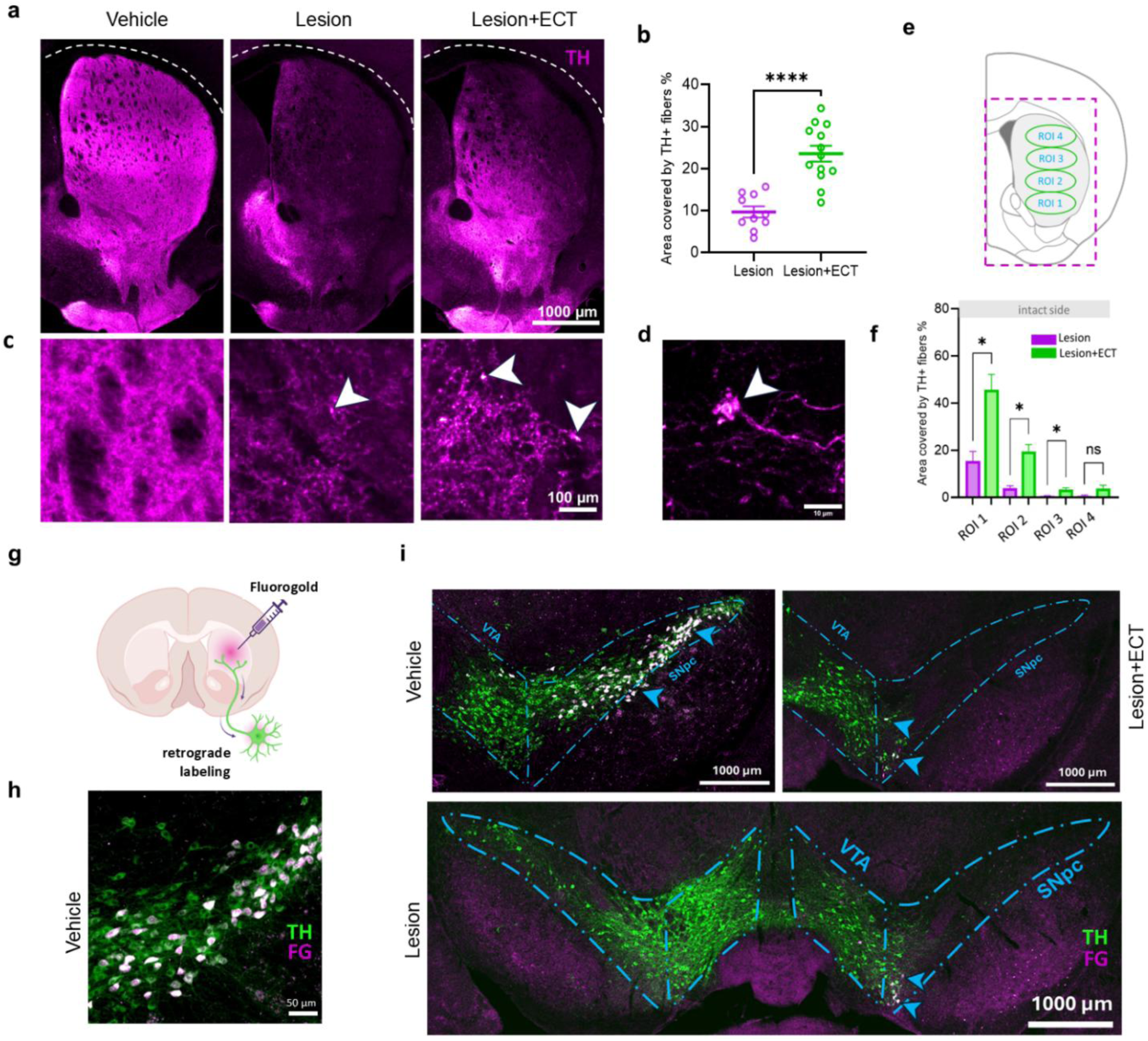
ECT promotes dopaminergic axon sprouting within the striatum. **(a)** Representative coronal striatal sections stained for tyrosine hydroxylase (TH) prepared at 3 days post intervention Scale bar, 1 mm. **(b)** Quantification of TH⁺ fiber density across the striatum, expressed as a percentage of the total area (Mann-Whitney test; Lesion, n=11; Lesion+ECT, n=12). **(c)** Magnified images of the dorsolateral striatum showing growth cone-like structures (arrowheads). Scale bar, 100 µm. **(d)** Image of a putative growth cone in the striatum of a Lesion+ECT mouse. Scale bar, 10 µm. **(e–f)** Regional quantification of TH⁺ fiber density in four striatal regions of interest (ROIs), from the most ventral (ROI 1) to the most dorsal (ROI 4). Significant reinnervation was observed in ventral ROIs (one-way ANOVA with post hoc Tukey’s test). **(g)** Experimental design for retrograde tracing: Fluorogold (FG) was injected into the dorsal striatum, and retrograde labeling was assessed in midbrain dopaminergic neurons 7 days post-injection. Created in BioRender.com. **(h–i)** Cells double-labeled for TH and FG appear white. In the intact striatum, FG labeling was predominantly observed in the lateral substantia nigra (SN) (h), whereas after 6-OHDA lesion, FG labeling shifted toward the border of the medial SN pars compacta (SNpc) and lateral ventral tegmental area (VTA) (i). No FG signal was detected in the contralateral hemisphere or medial VTA (Vehicle, Lesion, and Lesion+ECT, each n=1). *p < 0.05, ****p < 0.0001. Data represent mean ± SEM.

To further characterize the spatial distribution of reinnervation, the striatum was subdivided into four regions of interest (ROIs), spanning ventral to dorsal (**Fig. 2e**), as described before^34^. The most pronounced reinnervation was observed in the ventral striatum, suggesting preferential axonal outgrowth within that region (**Fig. 2f**).

To identify the origin of these reinnervating axons, we performed retrograde tracing by injecting Fluorogold (FG) into the lesioned medial-dorsal striatum using the same stereotaxic coordinates as for 6-OHDA injections (**Fig. 2g**). Seven days later, brains were processed for TH immunolabeling in the midbrain. In the intact brain, FG was primarily found in TH+ neurons of the lateral SN pars compacta (SNpc) (**Fig. 2h**; double-labeled cell bodies appear white). In contrast, in the lesioned hemisphere, FG labeling was observed in the medial SNpc and the adjacent lateral ventral tegmental area (VTA), indicating compensatory reorganization of nigrostriatal projections. No FG signal was detected in the contralateral hemisphere or medial VTA (**Fig. 2i**), consistent with the hypothesis that ECT promotes axonal sprouting from remaining neurons in the ipsilateral midbrain.

Together, these findings suggest that ECT promotes dopaminergic axonal regrowth within the lesioned striatum, a process that likely contributes to the observed motor recovery in treated animals.

### ECT reverses dopamine depletion-induced dSPN hyperexcitability

As dopamine depletion induces profound changes in the synaptic physiology of striatal SPNs^3,6,7^, we examined the effects of ECT on corticostriatal synapses and the intrinsic membrane properties of dSPNs and iSPNs. We focused our recordings on the dorsolateral striatum, which is implicated in motor control and is highly affected in PD. To distinguish dSPNs from iSPNs, D1-Cre or A2A-Cre mice were stereotaxically injected into the dorsolateral striatum with an adeno-associated virus encoding flox-tdTomato (**Fig. 3a-b**), concurrently with either saline or 6-OHDA injection.

**Figure 3.**
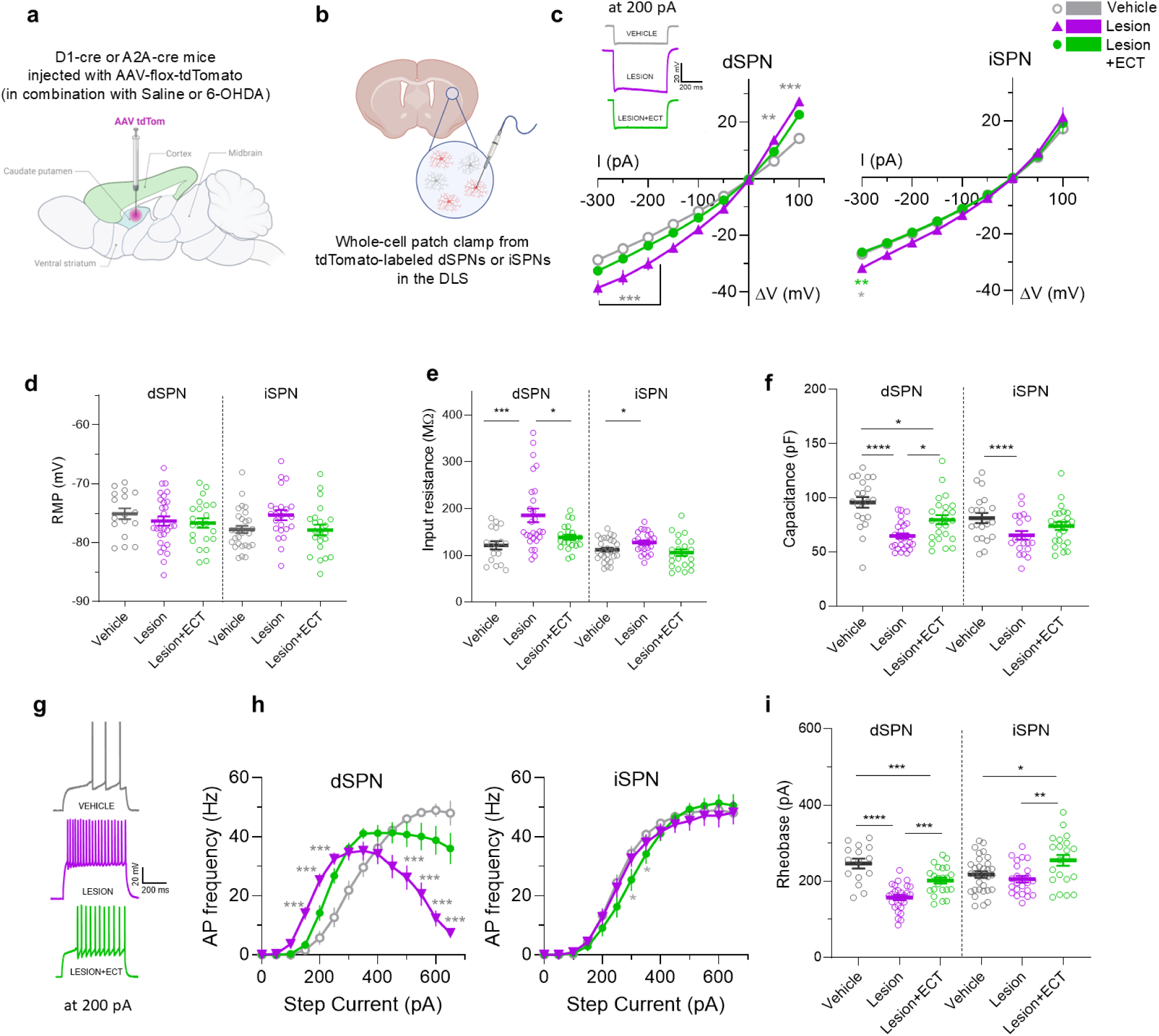
ECT normalizes lesion-induced alterations in intrinsic electrophysiological properties of striatal SPNs. **(a)** Strategy for Cre-dependent tdTomato labeling of dSPNs (D1-Cre) and iSPNs (A2A-Cre) using AAV-flox-tdTomato injections into the dorsolateral striatum. **(b)** Patch-clamp recordings were performed from fluorescently labeled SPNs in the dorsolateral striatum. The number of neurons (n) recorded from mice (N) was: dSPNs: Vehicle, n/N = 17/5, Lesion, n/N = 30/9, and Lesion+ECT, n/N = 22/8. iSPNs: Vehicle, n/N = 29/9, Lesion, n/N = 23/8, and Lesion+ECT, n/N = 22/7 mice). **(c)** In both types of SPNs, dopamine depletion led to a steeper I/V curve, indicating increased membrane conductance compared to vehicle-treated cells; these changes were reversed following ECT. **(d-f)** Effect of the dopamine lesion and ECT on cell resting membrane potential (d), input resistance (e) and membrane capacitance (f). **(g)** Representative traces of SPN responses to 200 pA. **(h)** Action potential spiking frequency in response to increasing current steps in dSPNs and iSPNs. **(i)** Rheobase (minimal current to elicit an action potential; measured by applying a current ramp) was reduced in dSPNs in the lesioned striatum and restored by ECT, suggesting normalization of intrinsic neuronal excitability. Plots show individual values with bars indicating mean ± SEM. One-Way ANOVA with Tukey’s post-hoc was used for group comparisons, Two-way ANOVA with Tukey’s post-hoc was used for curve comparisons. * p < 0.05, ** p < 0.01, *** p < 0.001, **** p < 0.0001. Panels a+b created in BioRender.com.

We first examined intrinsic electrophysiological cell properties, as dopamine depletion is known to induce compensatory homeostatic changes in SPN excitability^35,36^. Both dSPNs and iSPNs from dopamine-depleted animals exhibited increased (steeper) current-voltage (I/V) responses than the vehicle-treated group, while ECT treatment normalized the I/V curves to levels comparable to the vehicle group (**Fig. 3c**). While resting membrane potential remained unchanged, 6-OHDA lesions significantly increased input resistance and decreased membrane capacitance, changes that were normalized by ECT (**Fig. 3d-f**), indicating a reversal of the lesion-induced structural changes, including dendritic arborization and ion channel expression.

Next, we measured the intrinsic excitability of SPNs by applying 1 s somatic depolarizing step current to evoke trains of action potentials (APs) (**Fig. 3g**). In the dopamine-depleted striatum, dSPNs were hyperexcitable, exhibiting a leftward shift of the I/F curve and a reduced rheobase (the minimal current required to evoke an AP). ECT partially normalized both measures (**Fig. 3h-i**). In iSPNs, by contrast, ECT rendered neurons less excitable, reflected by an increased rheobase. Dopamine loss also caused spike-frequency adaptation, a progressive decrease in firing rate during sustained depolarization, specifically in dSPNs. This was reversed by ECT (**Fig. 3h**), likely reflecting restoration of D1 receptor-dependent hypoactivity in the parkinsonian state.

### ECT attenuates the increased glutamatergic corticostriatal drive in the parkinsonian striatum

Striatal dopamine depletion has been shown to alter glutamatergic inputs onto SPNs via both pre- and post-synaptic mechanisms^37^. Notably, our previous investigations^34^ demonstrated that dopaminergic axonal recovery, in turn, is regulated by glutamatergic signaling on dopamine growth cones, prompting us to investigate if ECT alters synaptic glutamatergic transmission in the striatum.

Measurements of miniature excitatory postsynaptic currents (mEPSCs) in the presence of a voltage-gated sodium channel blocker tetrodotoxin (TTX) confirmed that dopamine depletion increased both the frequency and amplitude of spontaneous vesicle fusion events in dSPNs and iSPNs, consistent with previous reports^38^. The increase in mEPSC frequency was fully reversed by ECT, indicating a presynaptic basis for this effect (**Fig. 4a-c**).

**Figure 4.**
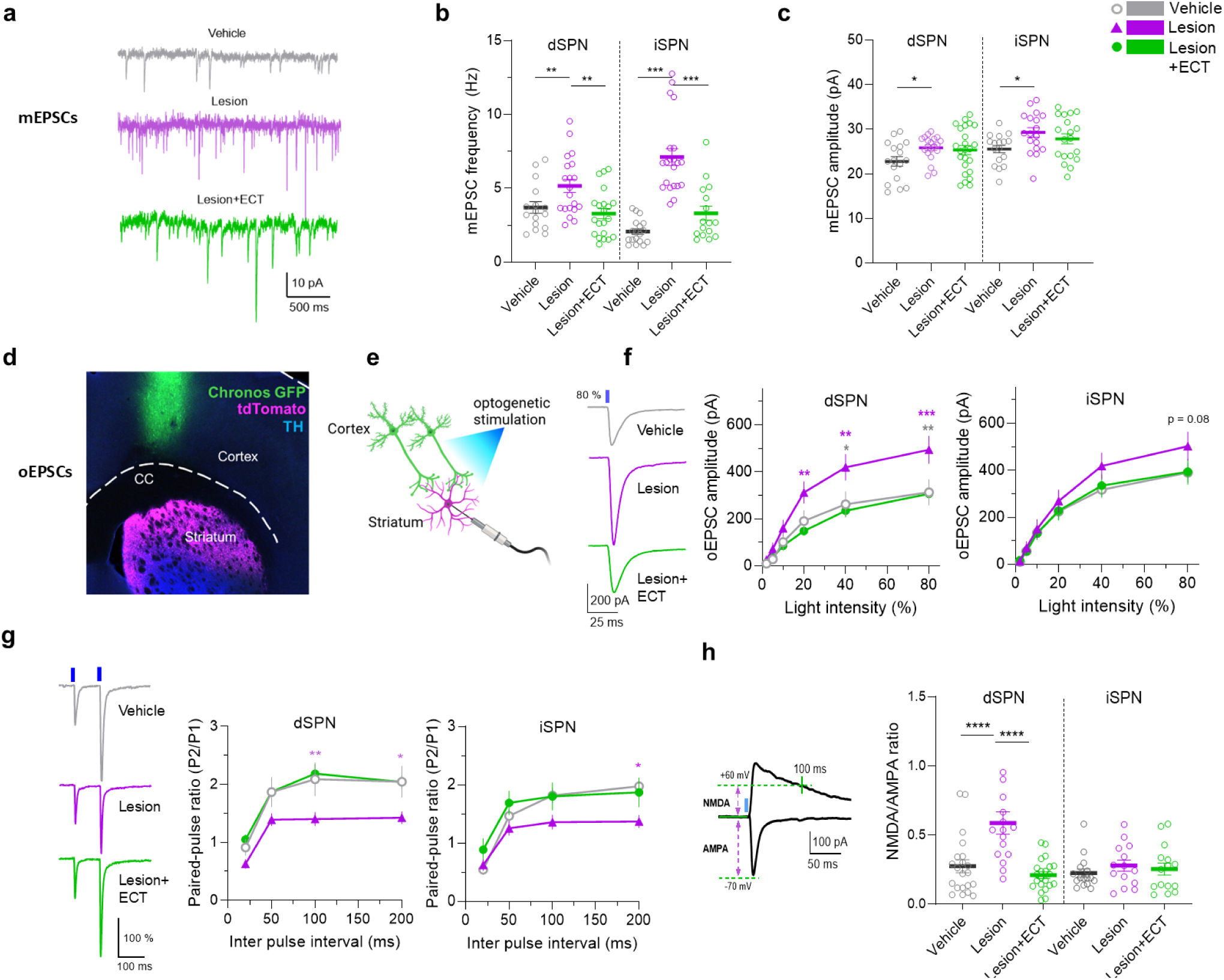
ECT restores lesion-induced changes at the corticostriatal synapse. **(a)** Representative traces showing spontaneous miniature excitatory postsynaptic currents (mEPSCs) recorded from dSPNs in the presence of 1 μM TTX. **(b-c)** 6-OHDA-lesioned mice exhibited significantly increased mEPSC frequency, indicating elevated presynaptic glutamatergic activity. These alterations were reversed by ECT. For mEPSCs, the number of neurons (n) recorded from mice (N) was: dSPNs: Vehicle n/N = 16/7, Lesion 20/8, Lesion+ECT 21/10. iSPNs: Vehicle n/N = 18/8, Lesion 21/8, and Lesion+ECT 16/7. **(d-e)** To enable pathway-specific recordings, tdTomato was selectively expressed in dSPNs or iSPNs in the dorsolateral striatum (as on Fig. 2a). Additionally, AAV expressing Chronos on synapsin promotor was expressed in the motor cortex neurons to allow optogenetic stimulation of corticostriatal projections. Panel e was created in BioRender.com. For oEPSCs, the number of neurons (n) recorded from mice (N) was: dSPNs: Vehicle, n/N = 21/7, Lesion 16/7, and Lesion+ECT 22/7. iSPNs: Vehicle, n/N = 14/8, Lesion 14/5, and Lesion+ECT 16/5). **(f)** Sample dSPNs traces of oEPSCs (left) and dependence of spike amplitudes on light stimulation intensity in dSPNs and iSPNs (right). **(g)** Representative traces of dSPNs paired-pulse responses at 100 ms inter-pulse interval (left, normalized to first EPSC) and dependence of paired-pulse ratio (PPR) on the inter pulse interval in different SPN types (right). **(h)** Representative oEPSC with indicated times to measure NMDA and AMPA EPSCs at + 60 and −70 mV respectively (left), and analysis of the NMDA/AMPA ratio (right). One-way ANOVA with post-hoc Tukey’s HSD. Curve comparisons (f+g) were analyzed by two-way ANOVA with post-hoc Tukey’s HSD. * p < 0.05, ** p < 0.01, *** p < 0.001, **** p<0.0001.

To further assess how ECT affects basal glutamatergic synaptic drive in the parkinsonian state, we virally expressed the channelrhodopsin Chronos^39^, in motor cortex neurons of D1 and A2A-tdTomato mice (**Fig. 4d**). Local optogenetic stimulation of corticostriatal afferents in the dorsolateral striatum (**Fig. 4e**) revealed that 6-OHDA lesioning increased the amplitude of optically evoked EPSCs (oEPSCs) in dSPNs, an effect that was reversed by ECT (**Fig. 4f**). In iSPNs, the increased oEPSC amplitudes did not reach significance.

It has been previously reported that corticostriatal presynaptic sites in the dopamine-depleted striatum show increased glutamate release probability, driven by synaptic fatigue and impaired D2 receptor-mediated presynaptic inhibition^40^. Consistent with this and Fig. 4b, dopamine depletion reduced the paired-pulse ratio (PPR) of oEPSCs, indicating a higher initial synaptic release probability. ECT fully reversed these changes in both dSPNs and iSPNs (**Fig. 4g**), suggesting that ECT can normalize the altered presynaptic plasticity induced by 6-OHDA.

To examine whether postsynaptic mechanisms are also affected by dopamine depletion and ECT, we measured NMDA and AMPA currents evoked by optogenetic activation of corticostriatal inputs. In agreement with previous reports^35^, dopamine depletion produced a marked increase in the NMDA/AMPA ratio in dSPNs, which was similarly normalized by ECT (**Fig. 4h**).

Together, these electrophysiological recordings indicate that most dopamine-depletion induced pre- and post-synaptic changes are fully or partially reversed by ECT, aligning with the behavioral recovery observed in 6-OHDA lesioned animals.

### ECT partially restores SPN morphology

To verify ECT-induced changes in striatal structural plasticity using an orthogonal assay, we examined SPN dendritic morphology in biocytin-loaded neurons. In dSPNs, which demonstrated a higher degree of 6-OHDA-induced plasticity, we also detected larger differences from the Sholl analysis of biocytin-labeled neurons^41,42^. Consistent with previous reports^35^, dopamine depletion caused a marked dendritic atrophy of dSPNs (**Fig. 5a**), reducing dendritic complexity and branching density as indicated by fewer dendritic intersections at each radius (**Fig. 5b-c**). We found that overall neuronal morphology, i.e., soma size or number of primary dendrites remained unchanged between treatment groups (**Fig. 5d**). Dendritic spine density, in contrast, was significantly reduced following 6-OHDA-lesion (**Fig. 5e-f**), in line with previous reports^43^. Notably, ECT treatment increased dendritic intersections at medium-length radii (50-90 µm in dSPNs or 50-70 µm in iSPNs), indicating enhanced arborization of secondary and tertiary dendrites. ECT thus normalized dopamine-depletion induced morphological deficits and restored dendritic complexity and spine density, consistent with our electrophysiological data.

**Figure 5.**
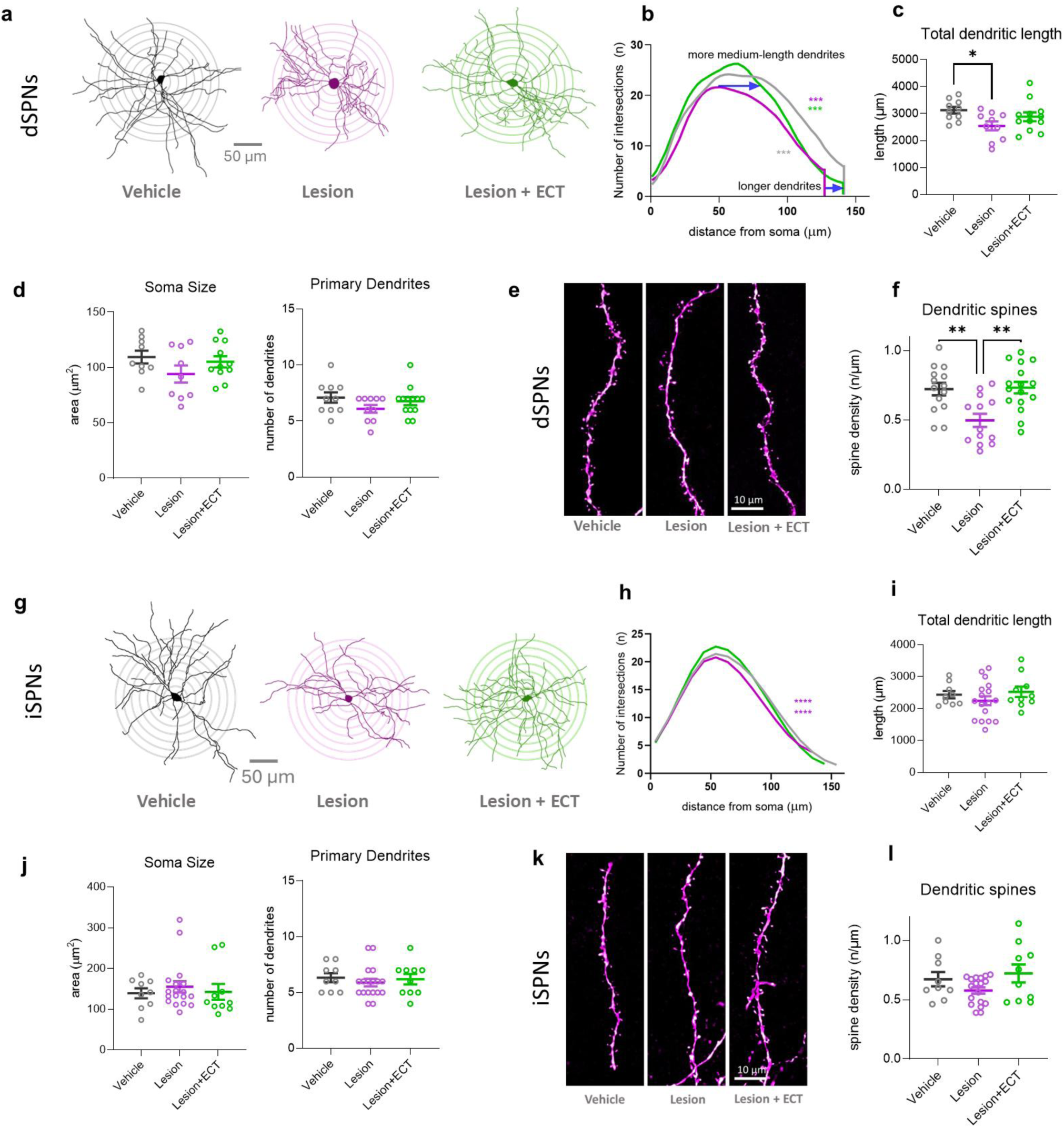
ECT restores lesion-induced alterations in striatal neuronal morphology. **(a)** Representative tracings of biocytin-filled direct pathway SPNs (dSPNs). **(b)** Sholl analysis of dendritic complexity showing the number of dendritic intersections at increasing distances from the soma. dSPNs from 6-OHDA-lesioned mice exhibited a reduction in medium-length dendrites (50–90 µm; two-way ANOVA at 10 µm intervals) and fewer long dendrites (≥130 µm), both of which were restored by ECT. **(c)** The total dendritic length of dSPNs was reduced following 6-OHDA lesioning. **(d)** Soma size and number of primary dendrites did not differ significantly across groups. **(e)** Representative images of dendritic segments from dSPNs from Vehicle, Lesion, and Lesion+ECT mice. **(f)** Dendritic spine density of dSPNs was reduced in lesioned mice and normalized by ECT (one-way ANOVA with post hoc Tukey’s HSD). **(g)** Representative tracings of biocytin-filled indirect pathway SPNs (iSPNs). **(h)** Sholl analysis of dendritic complexity in iSPNs revealed reduced dendritic branching in Lesion mice versus Vehicle controls, while Lesion+ECT mice displayed dendritic complexity comparable to Vehicle. Lesioned iSPNs showed fewer medium-length dendrites (70–100 µm), whereas ECT increased the number of short-to-medium dendrites (50–70 µm) (two-way ANOVA at 10 µm intervals). **(i–k)** Total dendritic length, soma size, and number of primary dendrites did not differ significantly between groups. **(l)** Representative images of dendritic spines from iSPNs from Vehicle, Lesion, and Lesion+ECT mice. **(m)** Reduction in dendritic spine density in distal dendritic segments of iSPNs following dopamine depletion was not statistically significant (one-way ANOVA). Biocytin-filled neurons (n) and mice (N): dSPNs: Vehicle n/N = 10/8, Lesion 10/9, Lesion+ECT 12/5. iSPNs: Vehicle n/N = 9/6, Lesion 18/7, and Lesion+ECT 10/5. *p < 0.05, **p < 0.01, ***p < 0.001. Data represent mean ± SEM.

Together, these results demonstrate that striatal SPNs, particularly dSPNs, become hyperexcitable and lose dendritic spines following dopamine depletion, and that ECT normalizes these effects. This normalization likely involves stabilization of glutamatergic synaptic activity. Interestingly, dopamine replacement therapy has been shown to alter SPN physiology, synaptic density, and corticostriatal connectivity in PD mouse models in an analogous fashion^44^, suggesting a potential convergence in the underlying mechanisms.

### ECT induces transcriptional reprogramming in dSPNs

To gain deeper insight into the molecular adaptations induced by ECT in the dopamine-depleted striatum, we performed single-nucleus RNA sequencing of striatal tissue from 6-OHDA-lesioned mice with and without ECT treatment. This revealed robust transcriptional responses to the procedure in several striatal cell types, including striatal SPNs. Here, we focus on transcriptional alterations in dSPNs.

Analysis of differentially expressed genes (DEGs) showed that ECT exerted effects on intracellular signaling pathways (**Fig. 6a, b**). *Pebp1*, an inhibitor of the Raf/MEK/ERK pathway, and *Camk2n1*, a CaMKII inhibitor, were strongly induced, possibly reflecting compensatory negative feedback mechanisms to heightened activity. The analysis further revealed strong upregulation of multiple components of mitochondrial oxidative phosphorylation, including *Cox6c*, *Cox4i1*, *Ndufs8*, *Cycs*, *Uqcr10*, *Atp5h*, and *Atp5g1 (***Fig. 6b**), consistent with a metabolic adaptation toward enhanced ATP production.

**Figure 6.**
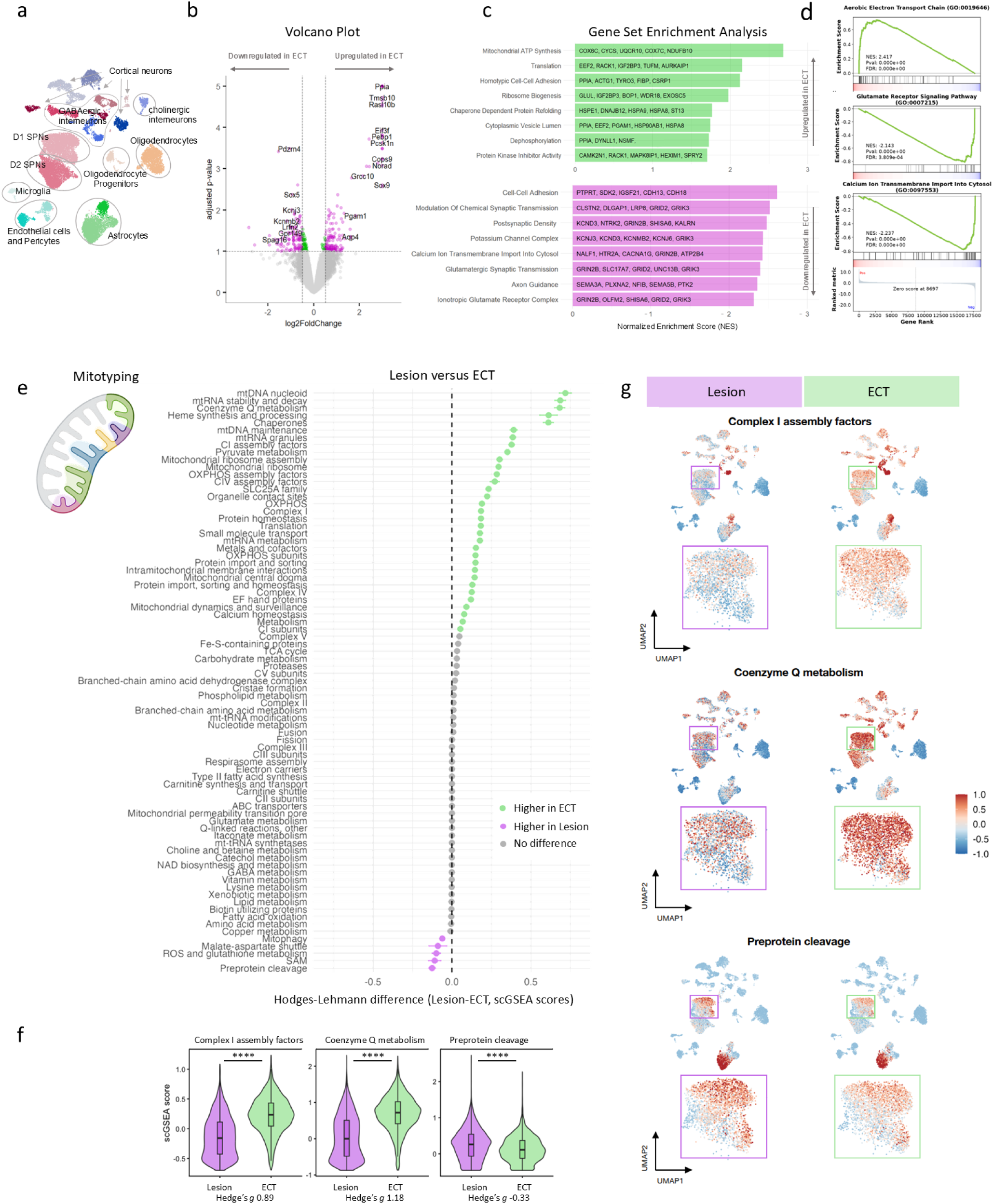
Transcriptomic changes in dSPNs following ECT in the dopamine-denervated striatum. **(a)** The dorsal striatum was dissected from 6-month-old male WT mice subjected to 6-OHDA lesion and sham treatment or ECT (n = 3 for Lesion or Lesion+ECT) and nuclei were isolated. CellTypist was used to cluster striatal cell types. Colors indicate cell clusters; distinct clusters reflect transcriptional heterogeneity. Only D1-expressing SPNs were selected for further analysis. **(b)** Volcano plot showing differentially expressed genes (DEGs) in D1-receptor-expressing SPNs. **(c)** GSEA of Upregulated and Downregulated Gene Sets: Bar plots showing the top significantly enriched gene sets (normalized enrichment score (NES) shown). **(d)** GSEA plots for selected enriched GO terms. Vertical, black lines in the middle indicate concentration of pathway genes at the positive/negative end of the gene rank, confirming significant enrichment (FDR <0.05, false discovery rate). Genes towards the left indicate upregulation in the Lesion+ECT group. **(e)** Mitotyping analysis using gene-to-pathway annotations from MitoCarta3.0^56^. Ranked Hodges-Lehmann estimates of pathway score differences, ordered from highest (higher in ECT relative to Lesion) to lowest (higher in Lesion relative to Sham). Pathways that met the inclusion criteria (absolute Hodges-Lehmann difference >0.05 z-score units and Benjamini-Hochberg adjusted p <0.05) are highlighted in color. Enriched programs included OxPhos assembly factors, Coenzyme Q metabolism, mtDNA/mtRNA maintenance. Reduced were preprotein cleavage pathways. **(f)** Violin plots of enrichment scores across dSPNs for select top pathways. **(g)** Feature plots for selected pathways. The outlined region highlights dSPNs. Enriched pathway expression is highlighted in darker shades of Red. ****p < 0.0001.

ECT further induced extensive cytoskeletal and structural remodeling as suggested by the upregulation of genes encoding tubulins (*Tuba1a*, *Tubb5*, *Tubb4b*), actins (*Actb*, *Actg1*), and neurofilaments (*Nefl*), consistent with dendritic or axonal reorganization^45^. Another upregulated gene is *Sox9*, which in neurons is associated with neurogenesis and synaptogenesis, specifically in SPNs^46^. Interestingly, *Nr4a2* (Nurr1), was upregulated in the lesioned striatum, and its downregulation by ECT may point to a reorganization of SPN identity or function in response to ECT. Nurr1 induction in the striatum has been found to depend on pathologically high activation of dSPNs in the parkinsonian state, and Nurr1 overexpression has been linked to altered SPN spine morphology and priming of levodopa-induced dyskinesia^47^. Conversely, key components of G-protein coupled signaling were downregulated, including *Gng5* and *Kcnj3* (GIRK1), suggesting altered synaptic responsiveness or excitability.

Gene set enrichment analysis (GSEA) revealed that ECT significantly downregulated gene sets associated with potassium channel function and glutamatergic signaling (**Fig. 6c,d**). GO terms encompassing glutamate receptor activity and synaptic transmission were suppressed, involving genes including *Slc17a7*, *Grid1*, *Grid2*, *Grik1*, *Grik2*, *Grik3*, *Grik4*, *Grik5*, *Grin2a*, *Gria3*, *Gria2*, and *Gria1*. These genes encode key components of AMPA, NMDA, and kainate receptors, as well as vesicular glutamate transporters and synaptic release machinery, and their repression is consistent with the attenuation of excitatory glutamatergic input and signaling within SPNs. Genes related to potassium channel activity, including *Kcnb1*, *Kcnc2*, *Kcnd3*, *Kcng1*, *Kcng2*, *Kcng3*, *Kcnh1*, *Kcnip4*, *Kcnj6*, *Kcnma1*, *Kcnmb2*, *Kcnq3*, *Kcnq5*, and *Kcns3* were also downregulated. These genes encode voltage-gated and G-protein-coupled inwardly rectifying potassium channels, essential for regulating neuronal excitability, action potential firing, and afterhyperpolarization. The suppression of excitatory signaling and ion channel activity after ECT may thus represent a restorative process aimed at rebalancing striatal output and preventing excitotoxic stress.

### ECT induces mitochondrial remodeling in dSPNs

Differential expression analysis revealed upregulation of mitochondrial oxidative phosphorylation (OxPhos)-related genes in dSPNs. GSEA further indicated metabolic remodeling involving a catabolic shift towards mitochondrial ATP synthesis, alongside an anabolic increase in protein translation and chaperone-mediated protein folding. Although GO terms such as *mitochondrial ATP synthesis* are commonly interpreted as reflecting a higher energy demand, the electron transport chain also plays a central role in maintaining redox balance and supporting biosynthesis processes required for cell and tissue maintenance^48^. Inhibition of the electron transport chain, and particularly Complex I, which is targeted by 6-OHDA^49,50^, impairs NADH oxidation to NAD+, leading to NADH accumulation and a shift toward “reductive stress” or mitochondrial redox imbalance^51,52^. Continuous electron flow through the electron transport chain is therefore essential not only to synthesize ATP demand, but also to regenerate NAD+, sustain tricarboxylic acid (TCA) cycle flux, and maintain mitochondrial membrane potential, whose collapse can trigger mitophagy^48,53^. Moreover, the TCA cycle supplies key biosynthetic intermediates and nucleotides necessary for anabolic growth^54^. Thus, the upregulation of OxPhos in response to ECT likely reflects a broader mitochondrial remodeling that extends beyond ATP synthesis.

To examine these non-ATP-related mitochondrial functions in dSPNs, we systematically profiled mitochondrial remodeling using a novel mitotyping pipeline^55^. Mitotyping leverages 149 MitoCarta 3.0-derived functional mitochondrial pathways^56^ to calculate pathway-level scores, capturing different aspects of mitochondrial biology. We applied single-cell GSEA across the entire dataset^55,57^, ranking enriched pathways (FDR<0.05) by the Hodges-Lehmann difference (Wilcoxon rank-sum test estimate) between ECT and Lesion groups. Consistent with the pseudobulk GSEA results, mitochondrial pathways in striatal dSPNs were mostly enriched in ECT compared to Lesion (**Fig. 6e**). The most highly upregulated MitoPathway with ECT was *mtDNA nucleoid*, the superstructure that encapsulates the mitochondrial genome^58^. Other robustly enriched programs included OxPhos assembly factors (**Fig. 6f,g**), coenzyme Q metabolism (a key electron carrier and ROS scavenger^59^), mtDNA and mtRNA maintenance pathways, mitochondrial chaperones (required for protein folding during biogenesis), and SLC25A family metabolite transporters linking mitochondrial and cytosolic metabolism. On the contrary, mitochondrial stress-related pathways such as ROS and glutathione metabolism and mitophagy showed modest reductions in ECT relative to Lesion (**Supplementary Fig. 2**). Pathways involved in preprotein cleavage (required for removal of mitochondrial targeting sequences (**Fig. 6g**) and protein sorting and assembly machinery) were also slightly reduced. These changes could indicate a potential bottleneck in mitochondrial protein import and processing. Collectively, these findings suggest that ECT triggers the coordinated remodeling of several mitochondrial programs, consistent with metabolic reorganization and potential induction of anabolic or biogenesis pathways, rather than a stress-associated response. Note that although we focused our analysis here on dSPNs, similar changes are observed in many other striatal cell populations, including iSPN, striatal GABA interneurons and astrocytes (**Fig. 6g, Supplementary Fig. 2**).

In summary, these findings indicate that ECT may enhance neuroplasticity through an upregulation of growth and metabolic pathways and activate protective mechanisms for recovery.

## DISCUSSION

We report that ECT exerts a comprehensive and long-lasting restorative effect on the parkinsonian striatum that includes enhanced dopamine axonal reinnervation. This regenerative response features dopamine-associated effects in the striatum, including a normalization of corticostriatal synaptic transmission, restoration of SPN morphology and electrophysiological properties, and recovery of both short- and long-term parkinsonian motor behaviors^7,35,60,61^.

The behavioral analysis demonstrates that ECT leads to robust and sustained improvement of a range of parkinsonian motor deficits. Lesioned mice treated with ECT exhibited increased overall motor activity, reduced motor asymmetry and recovery of physiological behaviors such as exploratory locomotion and rearing. These improvements reflect a restoration of more naturalistic and goal-directed behaviors typically disrupted by dopamine depletion. Mechanistically, this behavioral recovery can be attributed to two complementary processes: (1) dopaminergic reinnervation of the dorsal striatum, restoring modulatory input critical for balanced activity of direct and indirect pathway SPNs, and (2) structural and functional remodeling within the striatum, including normalization of SPN excitability, synaptic response and plasticity, and dendritic morphology.

The sustained behavioral recovery (>30 days post-ECT) is reminiscent of the “long-duration response” seen following L-DOPA (<10 days post L-DOPA), where sustained motor improvements are driven by synaptic plasticity^6,61^. However, chronic L-DOPA therapy has drawbacks in that its non-physiological, pulsatile stimulation eventually promotes the emergence of motor complications, notably L-DOPA-induced dyskinesias (LID), via maladaptive circuit remodeling^62,63^. Crucially, ECT achieved lasting recovery while simultaneously attenuating presynaptic corticostriatal glutamatergic tone and downregulating genes implicated in LID, specifically Grin2a (GluN2) and Kcnj6 (GIRK2)^63–65^. This suggests ECT may induce long-term neurorestoration via mechanisms that counteract homeostatic changes underlying LID.

Despite the extensive degeneration of SNpc dopaminergic neurons in PD, a subset of surviving neurons, particularly those located at the VTA-SNpc border, retains the intrinsic capacity for axonal regrowth^66^, as we have previously shown^34^. Here, we report that ECT stimulated these neurons to reinnervate the previously denervated striatum, restoring dopaminergic input critical for striatal function and motor behavior. Several factors likely contribute to the ability of surviving dopamine neurons to regenerate their axonal arbors. We find that ECT-induced neuronal activity and seizure-related excitation trigger gene expression programs associated with growth and plasticity, including cytoskeletal remodeling pathways. Second, metabolic and proteostatic effects appear to prime neurons for regeneration. Upregulation of OxPhos genes and protein homeostasis machinery provides the energetic and structural resources needed to support axonal growth. In addition, we had previously shown that balanced NMDA receptor signaling at growth cones is important for dopamine axonal sprouting^34,67^, and this process may be affected by ECT as indicated by alterations in glutamatergic cortico-striatal signaling and transcriptional shift in NMDA/AMPA receptor expression in striatal neurons.

Dopamine neurons are highly vulnerable to mitochondrial impairment due to their high metabolic demand, complex arborization, and pacemaking activity^68,69^ and mitochondrial dysfunction has also been observed in striatal SPNs following dopamine depletion^70,71^. This energetic deficit interferes with normal synaptic function and axonal maintenance and contributes to disease progression. An interplay between mitochondrial stress and proteostatic collapse contributes to a feedforward cycle of cellular dysfunction, ultimately disturbing neuronal integrity and synaptic plasticity in the striatum. The upregulation of mitochondrial and proteostasis-related pathway after ECT may reflect a neurorestorative response that enhances cellular resilience, promotes axonal maintenance, and supports recovery in the dopamine-depleted basal ganglia network. ECT may in part mimic physical exercise, in that it helps to maintain cellular homeostasis by “training” cellular bioenergetics and proteostasis^72^, resulting in adaptive remodeling, including upregulation of chaperones and mitochondrial biosynthesis.

Our mitotyping analysis suggests that ECT may influence mitochondrial organization and function on multiple levels. The coordinated upregulation of mtDNA-related pathways, mitochondrial protein translation, and chaperone activity, points to a restoration or rebuilding of mitochondrial components. Increased expression of OxPhos-related pathways could reflect greater energetic demand, but may also indicate efforts to re-establish redox balance, enhance TCA cycle flux efficiency, and support biosynthetic capacity for metabolite and nucleotide production during cellular repair. Together, these findings raise the possibility that improved mitochondrial performance contributes to enhanced cellular resilience. Complementary assays, such as mitochondrial density and enzyme activity^73^, could help to further clarify ECT’s impact on mitochondrial functional capacity.

Importantly, pathways downregulated by ECT, including those related to glutamatergic signaling and potassium channel function in dSPNs, are implicated in dopamine depletion-induced hyperexcitability, consistent with our electrophysiological findings showing normalization of SPN excitability following ECT. Chronic dopamine depletion leads to pathological remodeling of corticostriatal circuits, characterized by enhanced glutamatergic tone, increased ionic flux via NMDA receptors, and impaired synaptic and intrinsic regulation of firing in SPNs^36,74^. These changes may exacerbate excitotoxic stress in vulnerable neurons, particularly in dSPNs, which exhibit dendritic spine loss, increased NMDA/AMPA ratios, and reduced inhibitory regulation via GIRK and other K⁺ channels. A recent study using fosTRAP to characterize SPNs activated during levodopa-induced dyskinesias (LID) in a PD mouse model demonstrated a significantly increased presynaptic glutamatergic drive compared to fos-negative non-activated SPNs. The downregulation of excitatory receptor genes and potassium channels that we observe after ECT suggests that the treatment rebalanced intrinsic and synaptic excitability, potentially reducing excitotoxic stress and promoting a more stable physiological state within the striatal network. Given its efficacy in normalizing striatal hyperexcitability, it may be valuable to study the anti-dyskinetic effects of ECT.

Our study has several limitations. First, the acute 6-OHDA lesion model we employed does not recapitulate progressive α-synuclein PD pathology and it will be important to test ECT in models that more closely mirror the human disease. Second, we restricted our experiments to male mice, as 6-OHDA efficacy differs significantly between sexes, with females exhibiting estrogen-mediated neuroprotection^75,76^. Inclusion of female animals will be important in follow-up studies. Third, most of our molecular mechanistic data stems from dSPNs; future RNAseq analysis of midbrain dopaminergic neurons, which are most vulnerable in PD, are desirable, but could not be conducted in the present study due to the low number of surviving SNc dopamine neurons following lesion. Fourth, a critical consideration is whether this intervention, which resembles a state of eustress (beneficial stress), might have an upper limit. Although the sustained behavioral recovery observed in our model supports the overall beneficial nature of the treatment, it is unknown if long-term ECT becomes adverse by pushing cellular systems past their adaptive capacity. Future studies are warranted to implement and compare maintenance protocols (e.g., once weekly or once monthly ECT) with longer follow-up periods. Finally, clinical ECT protocols vary in electrode placement as well as in stimulation frequency and pulse width, and the optimal dosing parameters for promoting neurorestoration remain unknown; systematic preclinical investigations will be necessary to guide translational efforts. Moreover, ECT in mice affects the entire brain and likely penetrates deeper than in humans, where unilateral or focal current steering can be applied, potentially leading to differences in efficacy and side-effect profiles.

Together, our findings suggest that ECT promotes a neurorestorative environment capable of reestablishing functional corticostriatal circuits, ultimately supporting both the reemergence of physiological behaviors and the attenuation of PD-related motor impairments. The observed axonal sprouting and synaptic remodeling suggests that ECT reactivates developmental growth programs, as suggested by snRNA-seq upregulation of microtubule dynamics and mitochondrial biogenesis. The transcriptomic shift toward mitochondrial resilience and proteostasis suggests that ECT may counteract PD-related bioenergetic crisis.

As these improvements last a minimum of one month following treatment, in contrast to existing PD therapies that offer symptomatic relief, ECT appears to induce lasting neurorestoration by reversing key circuit-level dysfunctions with persistent motor improvements beyond the treatment period. Thus, our data aligns with clinical reports of ECT improving motor function in PD patients and might provide mechanistic underpinning for this observation^19^.

## CONCLUSION

PD provides a major therapeutic challenge, as current treatments are primarily symptomatic and fail to halt or reverse the underlying neurodegenerative processes. Dopamine replacement therapies and deep brain stimulation (DBS) offer significant symptomatic relief for many patients, although these do not attenuate further PD-related nigrostriatal degeneration^77,78^. In contrast, ECT, a noninvasive neurostimulation technique, has been reported to improve motor and psychiatric symptoms in PD, offering an attractive alternative, albeit the mechanism of ECT therapeutic effect has not been understood^79^. Our results position ECT as a neurorestorative therapy that repairs lost function through regeneration and plasticity and is distinct from symptomatic or neuroprotective interventions. As ECT is FDA-approved, its repurposing for PD could accelerate clinical translation, bypassing hurdles faced by novel pharmacologic therapies. Given its potential to target both motor and psychiatric symptoms, ECT could become a noninvasive, disease-modifying neurostimulation treatment for PD.

We note that while we have not measured effects of ECT on other neurotransmitter systems, similar effects of ECT on axonal outgrowth of monoaminergic neurons including those that release norepinephrine and serotonin, might underlie effects by which ECT treats depression.

## METHODS

### General experimental outline

A schematic of the experimental timeline and procedures is shown in **Fig. 1a**. A PD phenotype was induced by unilateral injections of the neurotoxin 6-hydroxydopamine (6-OHDA) into the striatum of wildtype C57BL/6 mice or transgenic animals expressing Cre in direct and indirect pathway spiny projection neurons (SPNs). Behavioral baseline data was obtained 3 weeks after 6-OHDA injection before the initiation of ECT or sham treatment. ECT or sham sessions were performed 10 times over the course of 2 weeks, and behavioral tests were repeated 3 days after the last treatment. Mice were then euthanized for electrophysiological recordings, immunolabeling, or RNA-seq. A separate cohort of mice underwent behavioral testing 30 days after the final treatment to assess long-term effects.

### Animals

The use of animals followed the National Institutes of Health guidelines and was approved by the Institutional Animal Care and Use Committee of Columbia University and New York State Psychiatric Institute (IACUC protocol #NYSPI-1680 and 1650). Animals were housed under standard conditions with free access to food and water under a 12:12-hour light-dark cycle (light 23:00-11:00 h).

As 6-OHDA toxicity differs significantly between sexes, we focused our experiments on male mice^75,76^. Adult C57BL/6 mice (3-6 months old; Jackson Laboratory #000664) were used for all experiments, except for differentiated morpho-physiological analyses of direct and indirect pathway SPNs, where we employed mice expressing Cre in D1 receptor-positive neurons (MMRRC_034258-UCD) or A2A receptor-positive neurons (MMRRC_036158-UCD).

### Dopamine neuron unilateral lesion

Mice received an intraperitoneal (i.p.) injection of desipramine (25 mg/kg, Tocris) to block norepinephrine transporters, thereby protecting noradrenergic neurons from 6-OHDA-induced toxicity and ensuring selective dopaminergic degeneration in the striatum and SN. Mice were next anesthetized with isoflurane and placed in a stereotaxic frame. A 6-OHDA hydrobromide (Tocris) solution (1.5 µl, 5 µg/µl, in 0.02% ascorbic acid in 0.9% saline) or saline alone (with 0.02% ascorbic acid) was infused at a rate of 0.2 µl/min for 10 min (total dose: 7.5 µg) through a glass micropipette inserted into the right striatum (coordinates relative to bregma: anteroposterior (AP), + 0.7; mediolateral (ML) −2.3; and dorsoventral (DV, coordinates relative to dura) −2.7 mm). 6-OHDA-lesioned mice received enhanced post-surgical care with fluid substitutions and high calorie food supplements (DietGel Boost, ClearH_2_O).

### Behavioral testing

Behavioral tests were performed three weeks after the 6-OHDA lesion during the awake cycle; mice were habituated to the behavioral room for at least 30 minutes before testing. To examine animal locomotion and exploratory activity, animals were tested in an Open Field arena (16 x 16 inches with 12-inch walls, modified without central light) for 20 minutes. The total distance traveled was recorded using tracking software (DeepLabCut^80^). To ensure the assessment of locomotor activity was independent of the confounding effects of habituation to a standard open-field box, we additionally measured the total distance traveled in a novel environment. This specialized environment was the *MoSeq* arena (see below for details), which consists of a circular enclosure with a sanded surface.

The severity of the unilateral motor impairment (lateralization) was assessed using the corridor test^29^. Mice were maintained on a restricted diet for 24 h to increase the motivation for sugar pellet retrievals. They were then placed in a narrow corridor with 10 pairs of small cups along the left and right walls containing five sugar pellets each. The number of sugar pellets retrieved and cup explorations on the left and right sides were counted by a blinded rater during a 7 minute interval. Lateralization ratio was calculated by dividing the sum of left retrievals and cup explorations by the sum of the total number of retrievals and cup explorations. Only mice with < 33% lateralization score were used for subsequent testing.

Motion sequencing (*MoSeq*^81^) was used to analyze spontaneous behavior in mice. Animals were individually placed in a circular open field arena, and their movements were recorded with a depth-sensing camera for 20 minutes. Behavioral data was processed using an unsupervised machine learning algorithm to identify and classify sub-second behavioral motifs (“syllables”), allowing quantitative assessment of treatment-induced changes in motor patterns. To determine the duration of the ECT effect, a separate cohort of mice underwent behavioral testing 30 days after the final treatment. Vehicle-treated control animals were included to account for nonspecific effects and natural variability over time.

### Electroconvulsive stimulation

After baseline behavior evaluation, mice were randomly assigned to receive either ECT or sham treatment of five interventions a week, ten sessions total. We performed trans-auricular ECT under isoflurane anesthesia (4.0 % induction, 1.0-2.0% maintenance) by attaching pre-wetted (0.9% saline) electrodes to both ears and delivering a brief electrical current (ECT Unit from Ugo Basile, 50 mA for 1 s, 100 Hz, 0.5 ms/pulse). The resulting tonic-clonic seizures lasted approximately 10 seconds (from seizure onset to muscle relaxation) and seizure onset and duration were monitored by tonic extension of the tail. Control mice were sham-treated with isoflurane anesthesia only (4.0 % induction, 1.0–2.0% maintenance over one minute).

### Brain slice preparation and *ex vivo* electrophysiology

To specifically label dSPN or iSPN populations, we injected a viral vector that expressed red fluorescent protein (250 nL AAV-FLEX-tdTomato, AddGene) into the dorsal striatum of D1-Cre and A2A-Cre mice at coordinates: AP +0.7, ML −2.3, DV −2.7. To optogenetically stimulate cortical axons in the striatum, an AAV encoding an ultrafast channelrhodopsin (250 nL AAV-Syn-Chronos-GFP, AddGene) was injected in the motor cortex at coordinates: AP +1.0, ML +1.0, DV −1.2). Viral injections were performed simultaneously to the 6-OHDA lesion.

Animals were euthanized by cervical dislocation 5 days after the last ECT or Sham treatment. Coronal 250 µm thick slices were prepared on a vibratome (VT1200; Leica, Germany) in oxygenated ice-cold cutting-artificial cerebrospinal fluid (ACSF) containing (in mM): 194 sucrose, 30 NaCl, 4.5 KCl, 26 NaHCO_3_, 6 MgCl_2_·6H_2_O, 1.2 NaH_2_PO_4_, and 10 D-glucose (pH 7.4, 290 ± 5 mOsm). Slices were collected and transferred to oxygenated normal ACSF containing (in mM) 125.2 NaCl, 2.5 KCl, 26 NaHCO_3_, 1.3 MgCl_2_·6 H_2_O, 2.4 CaCl_2_, 0.3 NaH_2_PO_4_, 0.3 KH_2_PO_4_, and 10 D-glucose (pH 7.4, 290 ± 5 mOsm) at 34°C and allowed to recover for at least 40 min before electrophysiological recordings. Slices were transferred to the recording chamber that was continuously perfused with normal ACSF (1.5-2 ml/min) at 34°C. Recording chamber was mounted on an upright differential interference contrast and microscope (Olympus BX50WI, Olympus, Tokyo, Japan) equipped with a 40x water immersion objective and an infrared video camera. Visualization of SPNs expressing tdTomato was achieved with 555 nm LED light illumination (Colibri 2, Zeiss, Germany) and DsRed filter cube. Patch pipettes (5-6 MΩ) were pulled using a PC-10 gravity puller (Narishige, Japan) and filled with pipette solutions as indicated below. Whole-cell patch clamp recordings were performed with a MultiClamp 700B amplifier (Molecular Devices, CA) and digitized at 10 kHz with InstruTECH ITC-18 (HEKA, Holliston, MA). Data was acquired using WinWCP software (University of Strathclyde, UK).

Intrinsic membrane properties and cell excitability were measured in whole-cell current clamp mode with pipette solution containing (in mM): 115 K-gluconate, 10 HEPES, 2 MgCl_2_6H_2_O, 20 KCl, 2 MgATP, 1 Na_2_-ATP, and 0.3 GTP (pH=7.3; 280 ± 5 mOsm). In voltage clamp mode, cells were held at −70 mV and optical excitatory postsynaptic currents (oEPSCs) were evoked by 470 nm LED light stimulation (5 ms) through a 40x objective lens at 10-80% of maximal light intensity controlled by a Master-9 (A.M.P.I., Jerusalem, Israel). The pipette solution contained (in mM): 120 CsMeSO_3_, 5 NaCl, 10 HEPES, 1.1 EGTA, 2 Mg^2+^-ATP, 0.3 Na-GTP, 2 Na-ATP, 10 TEA-4, and 5 QX-314 (pH=7.3, 280 ± 5 mOsm). A GABA_A_ receptor antagonist (picrotoxin, 25 µM) was bath-applied throughout recording to isolate glutamatergic transmissions. To isolate NMDA components, cells were held at +60 mV, and the NMDA EPSCs were measured at 100 ms following optogenetic stimulation. Miniature EPSCs were detected in the presence of 1 µM tetrodotoxin (TTX). Data was sampled at 20 kHz and filtered at 4 kHz. All chemicals for ACSF, pipette solution, and reagents were purchased from Sigma-Aldrich and Tocris.

### Biocytin filling and morphological analysis

SPNs were filled with biocytin (1% in the pipette solution) during whole-cell patch-clamp recordings, with one cell filled per slice. After recordings, slices were fixed in 4% paraformaldehyde (PFA) at 4°C for 24 hours and subsequently transferred to PBS. Following three PBS washes, slices were incubated in PBS containing 0.6% Triton X-100 (PBS-T) and streptavidin conjugated with AlexaFluor 647 or DyLight 649 (1:200) for 72 hours. Slices were then mounted, dried overnight, and coverslipped with anti-fading mounting medium (Fluoromount G). Images were acquired on a Leica SP8 confocal microscope using a 63x/1.4 NA Zeiss lens. For planar Sholl analysis, z-stacks were maximum-projected, and dendrites were manually traced to generate binary masks. Skeletonized masks were analyzed at 2 µm intervals using the SNT plugin in ImageJ^82^. Soma size, primary dendrites, and total branch points were manually counted by a blinded rater. For dendritic spine density analysis, high magnification images of dendritic segments were at 0.14 µm pixel resolution and 0.14 µm z-steps. Spine density was quantified on 2-4 dendritic segments per cell (mean segment length 25 ± 8 µm), located on average 87 ± 13 µm from the soma.

### Immunofluorescence and retrograde label

Animals were transcardially perfused with ice-cold phosphate-buffered saline (PBS) followed by 4% PFA. Brains were post-fixed in 4% PFA overnight. Coronal sections (50 micron) were collected serially using a vibratome (VT1200S, Leica) and processed for immunofluorescent labeling of tyrosine hydroxylase (TH), a key component of dopaminergic neurons. Free-floating sections were first blocked in PBS-T and 10% normal goat serum (NGS) for 1 h at room temperature. Next, they were incubated overnight in a mixture of primary antibodies against TH (1:500, Millipore AB9702) in PBS-T and 2% NGS at 4°C. The next day, after four 15 min washes in PBS-T, sections were incubated with fluorescent secondary antibodies (AlexaFluor 648, 1:1000, ThermoFisher) in PBS-T and 2% NGS at room temperature for 3h. Mounted slices were shortly dried, and a cover slip with an anti-fading mounting medium (Fluoromount G) was applied. Imaging was performed using a 10x air objective (Olympus), a corresponding fluorescence filter set and a 2.0 neutral density filter using an Orca Flash 4 V3 camera (Hamamatsu) and MetaMorph software (Molecular Devices) or cellSens (Olympus).

For retrograde tracing of striatal dopaminergic axons, animals of each condition (vehicle, lesion-sham, and lesion-ECT) received stereotaxic injections of 500 nl Fluorogold (FG, Fluorochrome) into the dorsal striatum, using the same coordinates as for the 6-OHDA lesion (relative to bregma: AP +0.7 mm; ML – 2.3 mm; and DV –2.7 mm relative to the dura). Seven days later, mice were transcardially perfused with PBS and PFA, brains sectioned and processed for immunofluorescent labeling of TH as described above. Imaging was conducted on a Leica SP8 confocal microscope; all imaging parameters were constant across experimental groups.

### Single-nucleus RNA sequencing (snRNA-Seq)

Mice were euthanized via cervical dislocation. The brain was removed from the skull, and the region of interest (right-hemispheric (lesioned) striatum) was quickly dissected on ice. Brains were placed on an ice-cold brain matrix (Zivic Instruments, Pittsburgh, PA) and separated into 1.0 mm sections using ice-cold razor blades. The striatum was dissected from slices between approximately +1.00 mm to 0.00 mm AP to Bregma. Tissue was flash frozen on liquid nitrogen and stored at −80°C.

All materials used were RNA-free, low-binding materials. Nuclei were extracted using Nuclei Extraction Buffer (Miltenyi, Germany) with Protector RNAse inhibitor (Sigma-Aldrich, final concentration 0.2 U/µl) with gentleMACS C tubes (Miltenyi). Samples were cooled with OctoMACS cooling elements during the dissociation on the gentleMACS Tissue Dissociator (program 4C_nuclei). Immediately after, the lysis buffer was quenched by adding 2 ml wash buffer to each tube. Nuclei were then processed as described previously (Starr Lab and ^83^). In short, nuclei were magnetically labeled with anti-nucleus microbeads (Miltenyi) and incubated for 15 minutes. The MACS separator was used to separate the labeled nuclei from cell debris (flow-through was discarded). In addition, the three samples per condition were hashed with anti-nuclear pore complex antibodies (Biolegend TotalSeq-B antibodies, mab414) and pooled to increase cost efficiency.

All samples were processed in-house at Columbia Genome Center core facility, which handled quality control, library construction, and sequencing to ensure consistency and reduce potential technical variability. We utilized RNA-seq with poly(A)-tail pulldown to enrich mRNAs from total RNA samples (STRDPOLYA), then proceeded with library construction using Illumina TruSeq chemistry. Libraries were then sequenced using Element AVITI at the Columbia Genome Center. Samples were multiplexed in each lane, which yielded a targeted number of paired-end 75bp reads for each sample. For converting BCL to FASTQ format, bases2fastq version 1.7.0.1196148384, coupled with adapter trimming was used. Pseudoalignment was performed to a Kallisto index created from transcriptomes (Mouse:GRCm39). The references and Kallisto version were updated on April 29, 2024 to Ensembl v111 and Kallisto 0.50.1. Sequencing was performed at a depth of 20 million reads per sample.

### snRNA-Seq Data Analysis

Striatal tissue samples from three biological replicates per experimental group were processed for single-nucleus RNA sequencing. Nuclei were isolated and hashed for multiplexing. Demultiplexing of pooled samples was performed using the DEMUX algorithm^84^ in conjunction with CellRanger and Loupe Browser (10x Genomics), applying sample-specific thresholds to ensure accurate assignment. On average, ∼6000 high-quality nuclei per donor were recovered for the striatum. Quality control steps included decontamination of ambient RNA with DecontX^85^, removal of nuclei with high mitochondrial or ribosomal gene content, filtering of lowly expressed genes, and normalization of library size. Data integration and clustering were carried out using the Phenograph-Leiden algorithm, with no batch effects observed across replicates. Cell type annotation was performed using CellTypist^86^, and clusters corresponding to D1-type and D2-type spiny projection neurons (dSPNs and iSPNs), as well as astrocytes, were selected for downstream analysis. Additionally, pseudogenes and unclassified mouse genes, often labeled with the prefix “Gm” or “Rik” were excluded to maintain biological relevance. Differential gene expression analysis was conducted using PyDESeq2^87^ after creating pseudobulks for each sample and cell type by summarizing the unnormalized counts after ambient RNA decontamination. Gene set enrichment analysis (GSEA) was performed using ClusterProfiler^88^ referencing *GO* gene ontology databases.

### snRNA-Seq mitotyping

Single-cell gene set enrichment analysis (scGSEA) was performed as described^55^ and executed in R version 4.3.0 on Ubuntu Linux using the gficf package version 2.0.0^57^. Raw counts (ambient RNA removed and rounded, mitochondrial DNA-encoded genes removed) were normalized using the gficf() function. scGSEA was performed with runScGSEA() using 50 random seeds to account for stochasticity, with MitoCarta gene-to-pathway annotations. Scores were z-scaled within each run across the dataset and then averaged across 50 runs. The averaged pathway scores were added to a Seurat object and visualized using the FeaturePlot() function. To statistically compare ECT vs Lesion, we used an unpaired Wilcoxon rank-sum test (two.sided) in R version 4.5.1 (macOS Sequoia 15.6). Effect sizes were computed using the Hodges Lehmann (HL) location estimator (ECT - Lesion), and 95% confidence intervals were obtained. P-values were adjusted for multiple testing using the Benjamini-Hochberg method. Pathways with an absolute HL difference > 0.05 z-score units and an adjusted p <0.05 were considered biologically shifted toward one condition. Positive values indicate higher enrichment in ECT, while negative values indicate higher enrichment in Lesion.

### Statistical analysis

Data was analyzed using pClamp (Molecular Devices, San Jose, CA), Igor Pro (Wavemetrics, Lake Oswego, OR), MATLAB (MathWorks, Natick, MA) and GraphPad Prism 8.0 (GraphPad Software Inc, San Diego, CA). Data are presented as mean ± standard error of the mean (SEM). Data sets with normal distributions were analyzed for significance using unpaired Student’s two-tailed t-test or analysis of variance (ANOVA) followed by Tukey’s post hoc test. Data sets with non-normal distributions were analyzed using the Mann-Whitney U test or Kruskal-Wallis test with Dunn’s adjustment for multiple comparisons. Data curves were analyzed with two-way ANOVA with Tukey’s post-hoc. The statistical tests are listed in the figure legends.

## ACKNOWLEDGEMENTS

The authors are grateful to the members of the Sulzer lab for their insight and discussions. We thank I. Serrano for her assistance in setting up the Motion Sequencing (MoSeq) equipment and V. Morales for her great help with animal husbandry and care during the research period. This project was supported by the Freedom Together Foundation (DS, AF) and NIH NIDA R0107418. AF and JB were supported by the Walter Benjamin Fellowship from the German Research Foundation (project numbers 504575882 and 523020841). This research used Genomics and High Throughput Screening Shared Resource and was funded in part through the NIH/NCI Cancer Center Support Grant P30CA013696.

## AUTHOR CONTRIBUTIONS

AF and DS conceptualized the study and designed the experimental framework. AF, JB, EVM, and DS interpreted the results. AF, SJC, JB, ACF and SB conducted data collection and analysis. ASM performed Mitotyping, and ASM and MP interpreted mitochondrial data. AF and DS drafted the manuscript. All authors reviewed and edited the manuscript.

---

**Supplementary Figure 1:**
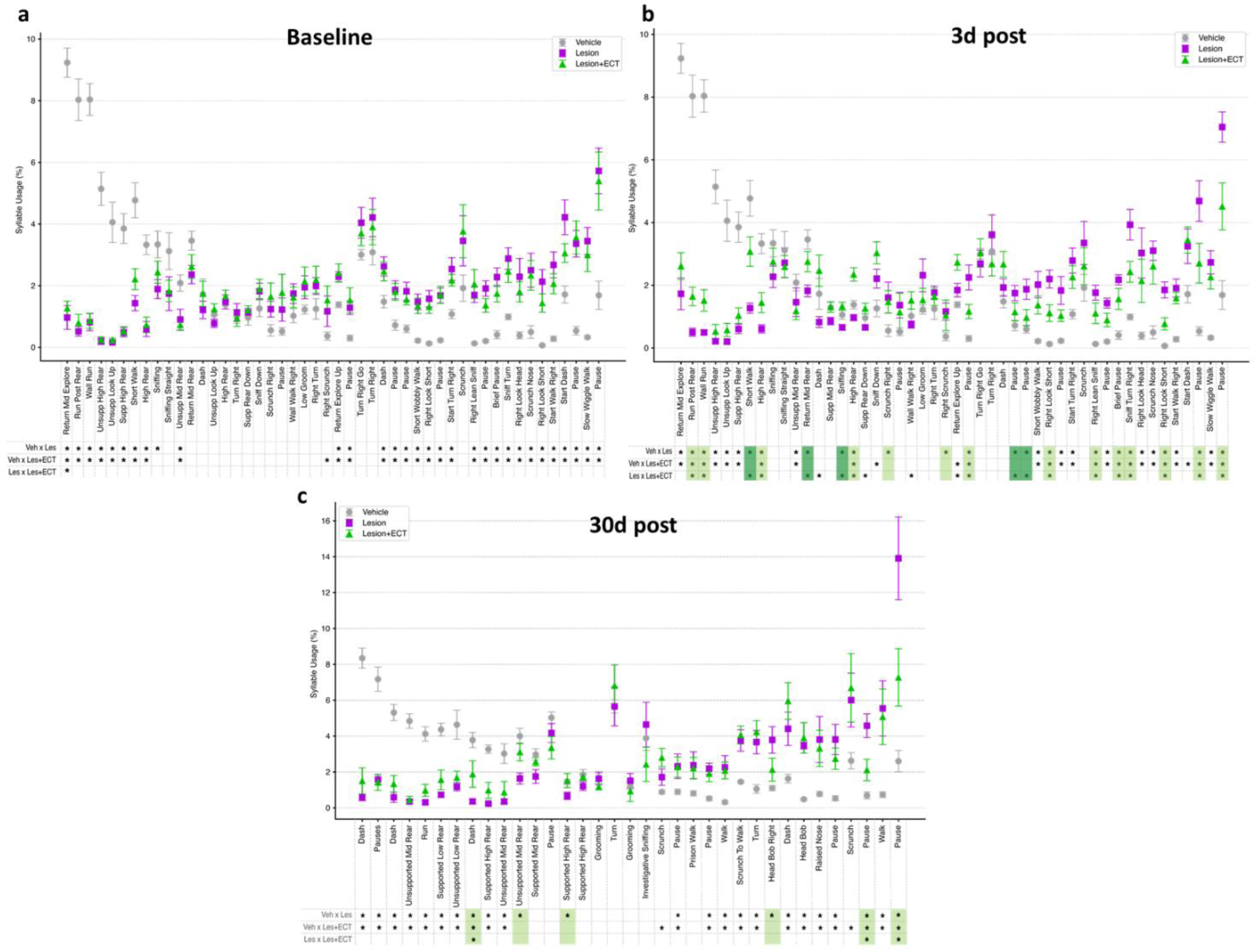
MoSeq Analysis of Mice Motor Performance Following ECT Intervention. MoSeq analysis of syllable (discrete, stereotyped units of movement) usage frequency at baseline **(a**, before intervention), three days **(b)** and thirty days **(c)** after ECT. Sample sizes at baseline and 3 days: vehicle (n=7), Lesion (n=11), and Lesion+ECT (n=13); at 30 days: vehicle (n=7), Lesion (n=5), and Lesion+ ECT (n=7). Randomization at baseline was effective, distributing animals with comparable PD-related motor behavior across each treatment group. Syllables are color-coded based on the treatment outcome: Dark green (Fully Reversed): significant difference between the Lesion and Lesion + ECT groups, and no significant difference between Vehicle and Lesion + ECT. Light green (Partially Reversed): significant difference between Lesion and Lesion + ECT, and significant difference remains between Vehicle and Lesion + ECT. Statistical analysis for comparisons across all three groups (Vehicle, ECT, Lesion) was performed using the Kruskal-Wallis test followed by post-hoc Dunn’s tests.

**Supplementary Figure 2:**
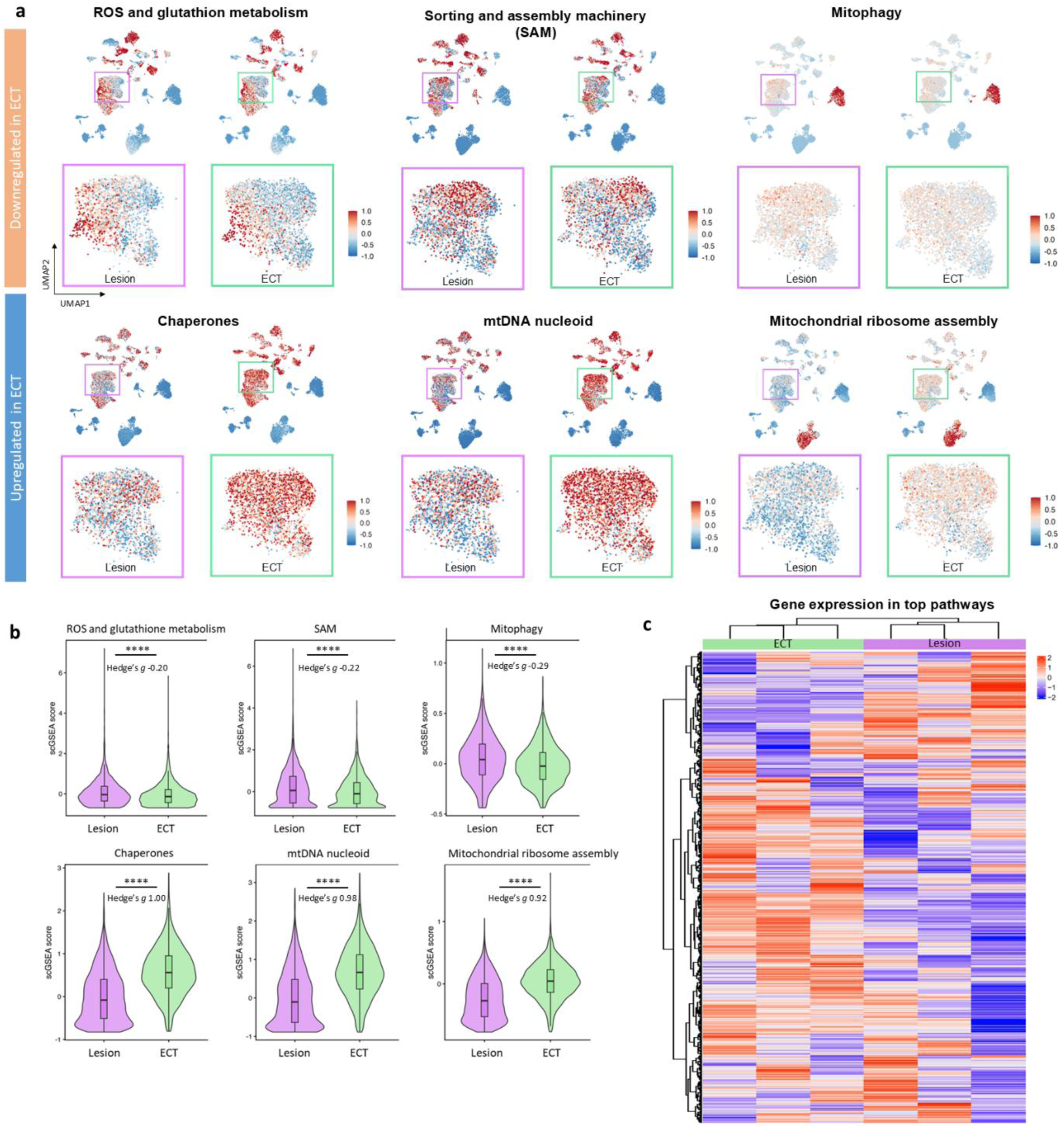
ECT Suppresses Mitochondrial Stress and Protein Processing Pathways in dSPNs. **(a)** Spatial feature plots showing the normalized expression score of genes related to the specific mitochondrial pathway. The outlined region highlights dSPN cluster identified by CellTypist. Downregulated mitochondrial pathways (upper row) showing negative enrichment score in Lesion+ECT relative to Lesion mice, including ROS (reactive oxygen species) and glutathione metabolism, SAM (sorting and assembly machinery) and mitophagy. Upregulated mitochondrial pathways (lower row), including mitochondrial chaperones, mtRNA nucleoid and mitochondrial ribosome assembly. **(b)** Violin plots of pathway enrichment scores of selected mitochondrial pathways. **(c)** Unsupervised hierarchical clustering heatmap and clustering of all mitochondrial genes included in the pathway calculations. The figure demonstrates successful separation and clustering of individual mice based on treatment (columns), highlighting the ECT-induced transcriptional shift. Genes are clustered by functional group (rows).

## REFERENCES

1. Kalia, L. V. & Lang, A. E. Parkinson’s disease. The Lancet 386, 896–912 (2015).

2. Arbuthnott, G. W., Ingham, C. A. & Wickens, J. R. Dopamine and synaptic plasticity in the neostriatum. Journal of Anatomy 196, 587–596 (2000).

3. Fieblinger, T. & Cenci, M. A. Zooming in on the small: The plasticity of striatal dendritic spines in l-DOPA–Induced dyskinesia. Movement Disorders 30, 484–493 (2015).

4. Haber, S. N. Corticostriatal circuitry. Dialogues in Clinical Neuroscience 18, 7–21 (2016).

5. Day, M. et al. Selective elimination of glutamatergic synapses on striatopallidal neurons in Parkinson disease models. Nat Neurosci 9, 251–259 (2006).

6. Nutt, J. G., Carter, J. H., Van Houten, L. & Woodward, W. R. Short- and long, duration responses to levodopa during the first year of levodopa therapy. Annals of Neurology 42, 349–355 (1997).

7. Witzig, V. S., Komnig, D. & Falkenburger, B. H. Changes in Striatal Medium Spiny Neuron Morphology Resulting from Dopamine Depletion Are Reversible. Cells 9, (2020).

8. Shen, W. et al. M4 Muscarinic Receptor Signaling Ameliorates Striatal Plasticity Deficits in Models of L-DOPA-Induced Dyskinesia. Neuron 88, 762–773 (2015).

9. Lang, A. E. & Espay, A. J. Disease Modification in Parkinson’s Disease: Current Approaches, Challenges, and Future Considerations. Movement Disorders 33, 660–677 (2018).

10. Kalia, L. V., Kalia, S. K. & Lang, A. E. Disease-modifying strategies for Parkinson’s disease. Movement Disorders 30, 1442–1450 (2015).

11. Rotheneichner, P. et al. Hippocampal neurogenesis and antidepressive therapy: Shocking relations. Neural Plasticity 2014, (2014).

12. Madsen, T. M. et al. Increased neurogenesis in a model of electroconvulsive therapy. Biological Psychiatry 47, 1043–1049 (2000).

13. Geddes, J. et al. Efficacy and safety of electroconvulsive therapy in depressive disorders: a systematic review and meta-analysis. The Lancet 361, 799–808 (2003).

14. Kellner, C. H., Obbels, J. & Sienaert, P. When to consider electroconvulsive therapy (ECT). Acta Psychiatrica Scandinavica 141, 304–315 (2020).

15. Luccarelli, J., Henry, M. E. & McCoy, T. H. Demographics of Patients Receiving Electroconvulsive Therapy Based on State-Mandated Reporting Data. J ECT 36, 229–233 (2020).

16. Weiss, A. et al. Royal Australian and New Zealand College of Psychiatrists professional practice guidelines for the administration of electroconvulsive therapy. Aust N Z J Psychiatry 53, 609–623 (2019).

17. Goldman, J. G. Non-motor Symptoms and Treatments in Parkinson’s Disease. Neurol Clin 43, 291–317 (2025).

18. Geduldig, E. T. & Kellner, C. H. Electroconvulsive Therapy in the Elderly: New Findings in Geriatric Depression. Curr Psychiatry Rep 18, 40 (2016).

19. Takamiya, A. et al. Electroconvulsive Therapy for Parkinson’s Disease: A Systematic Review and Meta-Analysis. Movement Disorders 36, 50–58 (2021).

20. Volkaerts, L., Roels, R. & Bouckaert, F. Motor Function Improvement after Electroconvulsive Therapy in a Parkinson’s Disease Patient with Deep Brain Stimulator. Journal of ECT 36, 66–68 (2020).

21. Fasano, A., Aquino, C. C., Krauss, J. K., Honey, C. R. & Bloem, B. R. Axial disability and deep brain stimulation in patients with Parkinson disease. Nat Rev Neurol 11, 98–110 (2015).

22. Andersen, K. et al. A double-blind evaluation of electroconvulsive therapy in Parkinson’s disease with “on-off” phenomena. Acta Neurologica Scandinavica 76, 191–199 (1987).

23. Laroy, M., Emsell, L., Vandenbulcke, M. & Bouckaert, F. Mapping electroconvulsive therapy induced neuroplasticity: Towards a multilevel understanding of the available clinical literature - A scoping review. Neurosci Biobehav Rev 173, 106143 (2025).

24. F, C., J, D., G, W., TM, M. & JR, Ng415. Sustained Ultrastructural Changes in Rat Hippocampal Formation After Repeated Electroconvulsive Seizures. The international journal of neuropsychopharmacology 23, 446–458 (2020).

25. Sartorius, A. et al. Correlations and discrepancies between serum and brain tissue levels of neurotrophins after electroconvulsive treatment in rats. Pharmacopsychiatry 42, 270–276 (2009).

26. Inta, D. et al. Electroconvulsive Therapy Induces Neurogenesis in Frontal Rat Brain Areas. PLoS ONE 8, 69869 (2013).

27. Alvarez-Fischer, D. et al. Characterization of the striatal 6-OHDA model of Parkinson’s disease in wild type and α-synuclein-deleted mice. Experimental Neurology 210, 182–193 (2008).

28. Slézia, A. et al. Behavioral, neural and ultrastructural alterations in a graded-dose 6-OHDA mouse model of early-stage Parkinson’s disease. Sci Rep 13, 19478 (2023).

29. Grealish, S., Mattsson, B., Draxler, P. & Björklund, A. Characterisation of behavioural and neurodegenerative changes induced by intranigral 6-hydroxydopamine lesions in a mouse model of Parkinson’s disease. European Journal of Neuroscience 31, 2266–2278 (2010).

30. Slézia, A. et al. Behavioral, neural and ultrastructural alterations in a graded-dose 6-OHDA mouse model of early-stage Parkinson’s disease. Sci Rep 13, 19478 (2023).

31. Lin, S. et al. Characterizing the structure of mouse behavior using Motion Sequencing. Nat Protoc 19, 3242–3291 (2024).

32. Dowd, E., Monville, C., Torres, E. M. & Dunnett, S. B. The Corridor Task: a simple test of lateralised response selection sensitive to unilateral dopamine deafferentation and graft-derived dopamine replacement in the striatum. Brain Res Bull 68, 24–30 (2005).

33. Carvalho, M. M. et al. Behavioral characterization of the 6-hydroxidopamine model of Parkinson’s disease and pharmacological rescuing of non-motor deficits. Mol Neurodegener 8, 14 (2013).

34. Schmitz, Y. et al. Glycine Transporter-1 Inhibition Promotes Striatal Axon Sprouting via NMDA Receptors in Dopamine Neurons. The Journal of Neuroscience 33, 16778 (2013).

35. Fieblinger, T. et al. Cell type-specific plasticity of striatal projection neurons in parkinsonism and L-DOPA-induced dyskinesia. Nat Commun 5, 5316 (2014).

36. Graves, S. M. & Surmeier, D. J. Delayed Spine Pruning of Direct Pathway Spiny Projection Neurons in a Mouse Model of Parkinson’s Disease. Front Cell Neurosci 13, 32 (2019).

37. Gardoni, F. & Bellone, C. Modulation of the glutamatergic transmission by Dopamine: a focus on Parkinson, Huntington and Addiction diseases. Front Cell Neurosci 9, 25 (2015).

38. Campanelli, F., Natale, G., Marino, G., Ghiglieri, V. & Calabresi, P. Striatal glutamatergic hyperactivity in Parkinson’s disease. Neurobiology of Disease 168, 105697 (2022).

39. Klapoetke, N. C. et al. Independent Optical Excitation of Distinct Neural Populations. Nat Methods 11, 338–346 (2014).

40. Bamford, N. S. et al. Dopamine Modulates Release from Corticostriatal Terminals. J Neurosci 24, 9541–9552 (2004).

41. Sholl, D. A. Dendritic organization in the neurons of the visual and motor cortices of the cat. J Anat 87, 387–406 (1953).

42. Horikawa, K. & Armstrong, W. E. A versatile means of intracellular labeling: injection of biocytin and its detection with avidin conjugates. Journal of Neuroscience Methods 25, 1–11 (1988).

43. Suarez, L. M., Solis, O., Aguado, C., Lujan, R. & Moratalla, R. L-DOPA Oppositely Regulates Synaptic Strength and Spine Morphology in D1 and D2 Striatal Projection Neurons in Dyskinesia. Cereb. Cortex 26, 4253–4264 (2016).

44. Nishijima, H., Ueno, T., Funamizu, Y., Ueno, S. & Tomiyama, M. Levodopa treatment and dendritic spine pathology. Movement Disorders 33, 877–888 (2018).

45. Maynard, Kristen. R., Hobbs, J. W., Rajpurohit, S. & Martinowich, K. Electroconvulsive seizures influence dendritic spine morphology and BDNF expression in a neuroendocrine model of depression. Brain stimulation 11, 856 (2018).

46. Song, X. et al. The SoxE factor Sox9 is selectively expressed in indirect pathway striatal projection neurons and regulates synaptogenesis. Fundamental Research https://doi.org/10.1016/j.fmre.2024.02.019 (2024) doi:10.1016/j.fmre.2024.02.019.

47. Sellnow, R. C. et al. Striatal Nurr1 Facilitates the Dyskinetic State and Exacerbates Levodopa-Induced Dyskinesia in a Rat Model of Parkinson’s Disease. J Neurosci 40, 3675–3691 (2020).

48. Monzel, A. S., Enríquez, J. A. & Picard, M. Multifaceted mitochondria: moving mitochondrial science beyond function and dysfunction. Nat Metab 5, 546–562 (2023).

49. Glinka, Y., Tipton, K. F. & Youdim, M. B. Nature of inhibition of mitochondrial respiratory complex I by 6-Hydroxydopamine. J Neurochem 66, 2004–2010 (1996).

50. Glinka, Y. Y. & Youdim, M. B. Inhibition of mitochondrial complexes I and IV by 6-hydroxydopamine. Eur J Pharmacol 292, 329–332 (1995).

51. Mick, E. et al. Distinct mitochondrial defects trigger the integrated stress response depending on the metabolic state of the cell. eLife 9, e49178 (2020).

52. Chen, W. W., Freinkman, E., Wang, T., Birsoy, K. & Sabatini, D. M. Absolute Quantification of Matrix Metabolites Reveals the Dynamics of Mitochondrial Metabolism. Cell 166, 1324–1337.e11 (2016).

53. Zorova, L. D. et al. Mitochondrial membrane potential. Anal Biochem 552, 50–59 (2018).

54. Inigo, M., Deja, S. & Burgess, S. C. Ins and Outs of the TCA Cycle: The Central Role of Anaplerosis. Annual Review of Nutrition 41, 19–47 (2021).

55. Monzel, A. S. et al. Mitotyping – An integrative framework to quantify mitochondrial specialization and plasticity. 2025.02.03.635951 Preprint at 10.1101/2025.02.03.635951 (2025).

56. Rath, S. et al. MitoCarta3.0: an updated mitochondrial proteome now with sub-organelle localization and pathway annotations. Nucleic Acids Res 49, D1541–D1547 (2021).

57 Franchini, M., Pellecchia, S., Viscido, G. & Gambardella, G. Single-cell gene set enrichment analysis and transfer learning for functional annotation of scRNA-seq data. NAR Genom Bioinform 5, lqad024 (2023).

58. Kukat, C. et al. Super-resolution microscopy reveals that mammalian mitochondrial nucleoids have a uniform size and frequently contain a single copy of mtDNA. Proc Natl Acad Sci U S A 108, 13534–13539 (2011).

59. Baschiera, E., Sorrentino, U., Calderan, C., Desbats, M. A. & Salviati, L. The multiple roles of coenzyme Q in cellular homeostasis and their relevance for the pathogenesis of coenzyme Q deficiency. Free Radic Biol Med 166, 277–286 (2021).

60. Paillé, V. et al. Distinct Levels of Dopamine Denervation Differentially Alter Striatal Synaptic Plasticity and NMDA Receptor Subunit Composition. J. Neurosci. 30, 14182–14193 (2010).

61. Barbato, L. et al. The long-duration action of levodopa may be due to a postsynaptic effect. Clin Neuropharmacol 20, 394–401 (1997).

62. Huang, Y.-T., Chen, Y.-W., Lin, T.-Y. & Chen, J.-C. Suppression of presynaptic corticostriatal glutamate activity attenuates L-dopa-induced dyskinesia in 6-OHDA-lesioned Parkinson’s disease mice. Neurobiology of Disease 193, 106452 (2024).

63. Campanelli, F., Natale, G., Marino, G., Ghiglieri, V. & Calabresi, P. Striatal glutamatergic hyperactivity in Parkinson’s disease. Neurobiology of Disease 168, 105697 (2022).

64. Kochoian, B. A., Bure, C. & Papa, S. M. Targeting Striatal Glutamate and Phosphodiesterases to Control L-DOPA-Induced Dyskinesia. Cells 12, 2754 (2023).

65. Yabumoto, T. et al. Striatal GluN2A gene suppression reduces L-DOPA-induced abnormal involuntary movements in parkinsonian rats. Neuropharmacology 279, 110616 (2025).

66. Fearnley, J. M. & Lees, A. J. Ageing and Parkinson’s disease: substantia nigra regional selectivity. Brain 114 (Pt 5), 2283–2301 (1991).

67. Schmitz, Y., Luccarelli, J., Kim, M., Wang, M. & Sulzer, D. Glutamate controls growth rate and branching of dopaminergic axons. Journal of Neuroscience 29, (2009).

68. Exner, N., Lutz, A. K., Haass, C. & Winklhofer, K. F. Mitochondrial dysfunction in Parkinson’s disease: molecular mechanisms and pathophysiological consequences. EMBO J 31, 3038–3062 (2012).

69. González-Rodríguez, P. et al. Disruption of mitochondrial complex I induces progressive parkinsonism. Nature 599, 650–656 (2021).

70. Schapira, A. H. Nuclear and mitochondrial genetics in Parkinson’s disease. J Med Genet 32, 411–414 (1995).

71. Chinta, S. J. & Andersen, J. K. Redox imbalance in Parkinson’s disease. Biochim Biophys Acta 1780, 1362–1367 (2008).

72. Gusdon, A. M. et al. Exercise Increases Mitochondrial Complex I Activity and DRP1 Expression in the Brains of Aged Mice. Exp Gerontol 90, 1–13 (2017).

73. Rosenberg, A. M. et al. Brain mitochondrial diversity and network organization predict anxiety-like behavior in male mice. Nat Commun 14, 4726 (2023).

74. Ryan, M. B., et al. Excessive firing of dyskinesia-associated striatal direct pathway neurons is gated by dopamine and excitatory synaptic input. Cell Reports 43, (2024).

75. Tamás, A., Lubics, A., Szalontay, L., Lengvári, I. & Reglodi, D. Age and gender differences in behavioral and morphological outcome after 6-hydroxydopamine-induced lesion of the substantia nigra in rats. Behav Brain Res 158, 221–229 (2005).

76. Santoro, M. et al. Mapping of catecholaminergic denervation, neurodegeneration, and inflammation in 6-OHDA-treated Parkinson’s disease mice. npj Parkinsons Dis. 11, 28 (2025).

77. Hilker, R. et al. Disease progression continues in patients with advanced Parkinson’s disease and effective subthalamic nucleus stimulation. *Journal of Neurology*, Neurosurgery and Psychiatry 76, 1217–1221 (2005).

78. Pal, G. D. et al. Comparison of neuropathology in Parkinson’s disease subjects with and without deep brain stimulation. Movement Disorders 32, 274–277 (2017).

79. Ruiz, P. J. G. Electroconvulsive Therapy and Movement Disorders. New Perspectives on A Time-Tested Therapy. Movement Disorders Clinical Practice 8, 521–524 (2021).

80. Mathis, A. et al. DeepLabCut: markerless pose estimation of user-defined body parts with deep learning. Nat Neurosci 21, 1281–1289 (2018).

81. Lin, S. et al. Characterizing the structure of mouse behavior using Motion Sequencing. Nat Protoc 19, 3242–3291 (2024).

82. Arshadi, C., Günther, U., Eddison, M., Harrington, K. I. S. & Ferreira, T. A. SNT: a unifying toolbox for quantification of neuronal anatomy. Nat Methods 18, 374–377 (2021).

83. Gaublomme, J. T. et al. Nuclei multiplexing with barcoded antibodies for single-nucleus genomics. Nat Commun 10, 2907 (2019).

84. Zhu, Q., Conrad, D. N. & Gartner, Z. J. deMULTIplex2: robust sample demultiplexing for scRNA-seq. Genome Biology 25, 37 (2024).

85. Yang, S. et al. Decontamination of ambient RNA in single-cell RNA-seq with DecontX. Genome Biol 21, 57 (2020).

86. Xu, C. et al. Automatic cell-type harmonization and integration across Human Cell Atlas datasets. Cell 186, 5876–5891.e20 (2023).

87. Love, M. I., Huber, W. & Anders, S. Moderated estimation of fold change and dispersion for RNA-seq data with DESeq2. Genome Biology 15, 550 (2014).

88. Yu, G., Wang, L.-G., Han, Y. & He, Q.-Y. clusterProfiler: an R Package for Comparing Biological Themes Among Gene Clusters. OMICS 16, 284–287 (2012).

